# Developmental Emergence of Replay–DMN Coordination Predicts Entorhinal Grid Codes

**DOI:** 10.1101/2025.04.02.646732

**Authors:** Luoyao Pang, Yukun Qu, Jianxin Ou, Shuqi Wu, Yuejia Luo, Chetan Gohil, Mark Woolrich, Tim Behrens, Yunzhe Liu

## Abstract

Neural replay, the reactivation of experience-related activity patterns during offline periods, has been implicated in organising memories into cognitive maps. However, how replay processes mature through childhood and adolescence remains unclear. Here we studied 106 participants aged 8–25 who learned a hidden two-dimensional (2D) associative structure and then underwent magnetoencephalography (MEG) during a map-based inference task and post-task rest to detect spontaneous replay. We found that during post-task rest, younger individuals relied primarily on replay alone to represent the 2D relationships, whereas older individuals showed increasingly precise timing alignment between replay events and activity in the default mode network (DMN), particularly when replay occurred at the trough of a DMN theta oscillation. Such replay–DMN coupling predicted the strength of a grid-cell-like code in the entorhinal cortex identified in this cohort with fMRI. Furthermore, resting-state fMRI revealed that DMN functional connectivity strengthened with age, in parallel with reduced reliance on replay and more efficient inference performance. These results reveal a developmental progression in which replay shifts from an isolated process in youth to a coordinated DMN mechanism by adulthood. This transition predicts the maturation of grid-like schema representations, shedding light on how children gradually build internal knowledge structures to support flexible inference.

## Introduction

Neural replay, the reactivation of neural patterns reflecting past experiences during rest or sleep, is a well-established mechanism for memory consolidation and is classically associated with hippocampal sharp-wave ripples (Carr et al., 2011; Diba & Buzsáki, 2007; Skaggs & McNaughton, 1996). Replay has also been linked to flexible inference in humans (Liu et al., 2019; Schwartenbeck et al., 2023). Rodent studies show that replay is present even before weaning and becomes progressively more extended and structured with maturation (Farooq & Dragoi, 2019; Muessig et al., 2019). However, the developmental trajectory of replay in humans, and its specific role in cognitive development, remain largely unexplored.

Replay has been proposed to help construct internal representations that guide flexible behaviour (Behrens et al., 2018; Whittington et al., 2022). It may do so either by forming associative maps via local links between related experiences, such as a successor representation (Barron et al., 2020; Stachenfeld et al., 2017), or by integrating new information into pre-existing schemas—abstract knowledge structures (Piaget, 1952; Tolman, 1948) that support inferences beyond direct experiences (Ou et al., 2025; Xiao et al., 2025). Consistent with this idea, a recent developmental fMRI study of 203 participants (ages 8–25) reported that inference ability improves with age in parallel with increasing expression of a schema-like representation (Qu et al., 2025), manifested as grid-cell-like coding in the entorhinal cortex (EC) (Hafting et al., 2005; Killian et al., 2012). In younger individuals who had not yet formed a clear grid-based schema, the hippocampus appeared to play a larger compensatory role when performing inference task, suggesting a heavier reliance on hippocampal mechanisms before a shift toward a grid-like schema representation. Such grid-like coding has been observed in both physical and conceptual spaces in the human brain (Constantinescu et al., 2016; Doeller et al., 2010; Park et al., 2021).These findings raise the intriguing question of whether neural replay underlies this developmental transition and how it might relate to the emergence of entorhinal grid codes.

Addressing this question requires considering the coordination of replay with broader cortical networks. In adults, replay events often coincide with activation of the default mode network (DMN), and are accompanied by hippocampal sharp-wave ripples (Higgins et al., 2021; Norman et al., 2021; Vaz et al., 2020). Notably, the DMN undergoes substantial maturation during childhood and adolescence (Fair et al., 2008; Fan et al., 2021), with enhanced connectivity in key hubs such as the posterior cingulate cortex (PCC) (Supekar et al., 2010). It also displays increased specialisation and integration across regions, including privileged links with the hippocampus (Ezama et al., 2021; Olsen et al., 2017; Ward et al., 2014) and stronger internal connections (Fair et al., 2009; Supekar et al., 2010). These developmental changes suggest that replay might gradually come to be coordinated with DMN activity as the brain network architecture matures.

Here, we investigated how replay activity and its coordination with the DMN evolve from childhood to early adulthood, and whether this coordination supports the development of entorhinal grid-like codes. To this end, 106 participants aged 8–25 were trained on a hidden 2D rank-order structure and then performed a map-based inference task, followed by a post-task rest period during MEG recording. This design enabled us to detect spontaneous replay events after learning and to track transient DMN activation states. We also tested whether the precise timing of replay relative to ongoing DMN activity would predict each individual’s entorhinal grid coding strength, measured previously with fMRI in the same cohort (Qu et al., 2025).

Consistent with our hypothesis, we found a clear developmental shift: younger participants exhibited more frequent replay events in isolation, whereas older participants showed more selective replay events tightly coordinated with DMN activity, particularly when replay coincided with the trough of a theta-band DMN oscillation. This replay–DMN alignment in older individuals was associated with stronger DMN co-activation during replay and with the presence of more robust grid-like representations in the EC. These results suggest that as the brain matures, replay transitions from an isolated mechanism to a system-level process coordinated with DMN network oscillations, potentially facilitating the formation of schema-like internal maps for flexible inference.

## Results

### Age-related improvement in map-based inference

All participants underwent extensive off-scan behavioural training, before MEG scanning (**Figure 1A**; see Methods for details and Qu et al., 2025). During this training, they learned the adjacent rank order (differing by one rank) of 25 objects for attack and defence power, corresponding to the two dimensions of a hidden 5 × 5 2D grid. Through repeated pairwise comparisons along each dimension, participants gradually acquired the power hierarchy of the objects. Only those who reached at least 80% accuracy on all pairwise relationships proceeded to the subsequent experiment. This thorough training ensured that participants of all ages fully understood the task, minimised fatigue during scanning, and controlled for other age-related confounds (see also **Supplementary Note 1**).

**Figure 1.**
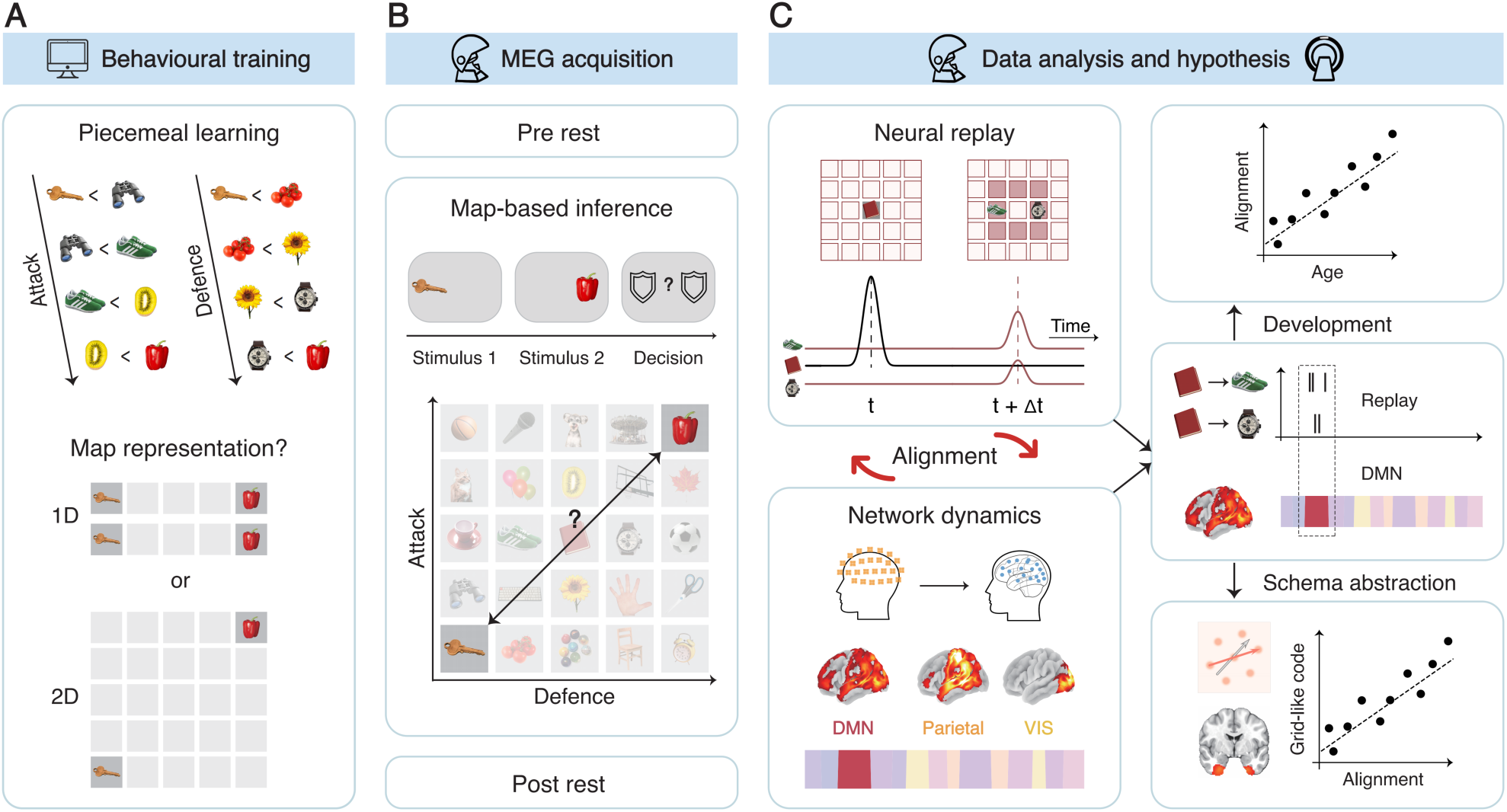
Overview of experimental design and analysis. (**A**) Behavioural training. Prior to scanning, participants completed an off-scan training protocol in which they memorised pairwise associations between 25 objects based on their attack or defence power. Participants were unaware of the underlying 5 × 5 2D structure and believed they were performing a simple memory task. Only those achieving at least 80% accuracy on all pairwise relationships proceeded to the neuroimaging sessions (see also Methods). (**B**) MEG acquisition. Participants first underwent a five-minute resting-state scan (eyes open). They then performed a map-based inference task, in which each trial presented two objects (1.5 s each) separated by a 0.4–0.6 s interval. After the second object, a decision cue indicated the dimension (attack or defence) to be compared, and participants had up to 2 s to choose the object with the higher rank. No feedback was given. This inference task was immediately followed by another five-minute resting-state scan, again with eyes open. (**C**) Data analysis and hypothesis. MEG recordings from the resting-state scans were used to quantify neural replay of experiences and its relationship with dynamic activity across different resting-state networks, with a focus on the DMN. We investigated whether replay events aligned with DMN activation and whether this alignment changed with development. We further assessed the functional relevance of replay–DMN alignment by testing its relationship to grid-like code in the EC measured in fMRI (Qu et al., 2025).

Following this training, participants performed an inference task and completed pre- and post-task resting-state scans under MEG (**Figure 1B**). The pre-task rest occurred before the inference task, and the post-task rest immediately after. These sessions enabled us to assess how the inference task affected neural replay during rest and how replay changed with age. After the MEG session, participants underwent fMRI scanning, which included a resting-state fMRI for linking with MEG replay measures, followed by an fMRI-scanned inference task reported in detail in Qu et al. (2025), where a robust grid-like code in the EC was identified.

In the MEG-scanned inference task, participants were presented with two randomly selected objects and asked to infer which had higher power in a cued dimension (attack or defence). Each object appeared on screen for 1.5 s, separated by a 0.4–0.6 s interval. After both objects were shown, a decision cue indicated which dimension to compare, and participants had up to 2 s to select the object with the higher rank on that dimension (**Figure 1B**). This procedure required them to make inferences beyond the pairs directly learned during training, potentially relying on an internal 2D map of the task space.

In total, 106 participants aged 8–25 years took part in the MEG experiment. After excluding those with excessive head movement, 98 participants (mean ± SEM age: 16.40 ± 0.64 years) remained in the final analysis (see **Supplementary Table 1**). Behaviourally, inference accuracy positively correlated with age (*r* = 0.219, *p* = 0.030), indicating that older participants made more correct inferences, and reaction time negatively correlated with age (*r* = −0.259, *p* = 0.010), suggesting faster responses among older participants. Notably, memory accuracy after training was not significantly associated with age (*r* = 0.074, *p* = 0.471), and the correlations between age and both accuracy and reaction time in the inference task remained significant when controlling for memory accuracy. The Advanced Progressive Matrices (APM) test, a standard IQ measure of reasoning, also improved significantly with age (*r* = 0.505, *p* < 0.001), and was significantly associated with inference performances (see **Supplementary Figure 1B**). These results demonstrate that as individuals mature, they become both more accurate and more efficient at complex inferences, in line with typical age-related gains in reasoning skills.

These improvements in inference ability may emerge as the 2D schema representation in the EC becomes more robust (Qu et al., 2025). However, it remains unclear whether the neural processes supporting this form of advanced knowledge organisation also mature with age. We hypothesised that spontaneous neural replay during rest is crucial to this development. In the sections that follow, we examine whether and how replay transitions across ages, including its coordination with the DMN (**Figure 1C**).

### Stronger replay for representing 2D map in younger individuals

We next examined neural replay patterns across ages. We first validated that the neural recording was not confounded by age-related factors. In developmental neuroscience studies such as this, potential confounds can arise from smaller head size in younger subjects (Brookes et al., 2018) or greater head movement in children (Wehner et al., 2008). For instance, smaller heads place the brain farther from the sensors, lowering the signal-to-noise ratio (SNR), while excessive movement degrades signal quality and introduces noise, along with other age-related factors. To address these issues, we evaluated the effects of head size and head movement on signal strength across our cohort (see Methods for details) and found they did not pose significant problems (see **Supplementary Note 1**). Moreover, we accounted for head size and head movement in all subsequent neural analyses to ensure the robustness of our results.

The replay detection pipeline begins with the neural decoding of stimuli during their visual presentation in the inference task (**Figure 2A**). Specifically, we trained object-specific decoders using task-evoked MEG data from all planar channels. For example, the decoder for the target object “book” showed higher predicted probabilities for “book” than for other decoders, indicating successful identification (**Figure 2B**). Decoding accuracy was highest when training and testing occurred at the same time point (**Figure 2C**). Across participants, cross-validated decoding accuracy peaked at around 300 ms after stimulus onset (one-sample *t*-test vs. chance, *t*(97) = 12.622, *p* < 0.001; **Figure 2D**). No significant correlation was observed between participant age and the timing of this peak (*r* = −0.176, *p* = 0.08). We therefore trained and tested decoders at the 300 ms peak for all subsequent analyses and for all subjects. Decoding accuracy at 300 ms was positively correlated with age (*r* = 0.255, *p* = 0.011; **Figure 2E**), and this relationship remained significant when controlling for head motion (*r* = 0.220, *p* = 0.043), head size (*r* = 0.242, *p* = 0.016), memory (*r* = 0.245, *p* = 0.016) and task (*r* = 0.202, *p* = 0.047) performances. These findings suggest that older participants demonstrated more distinct and robust neural representations of the objects during the task.

**Figure 2.**
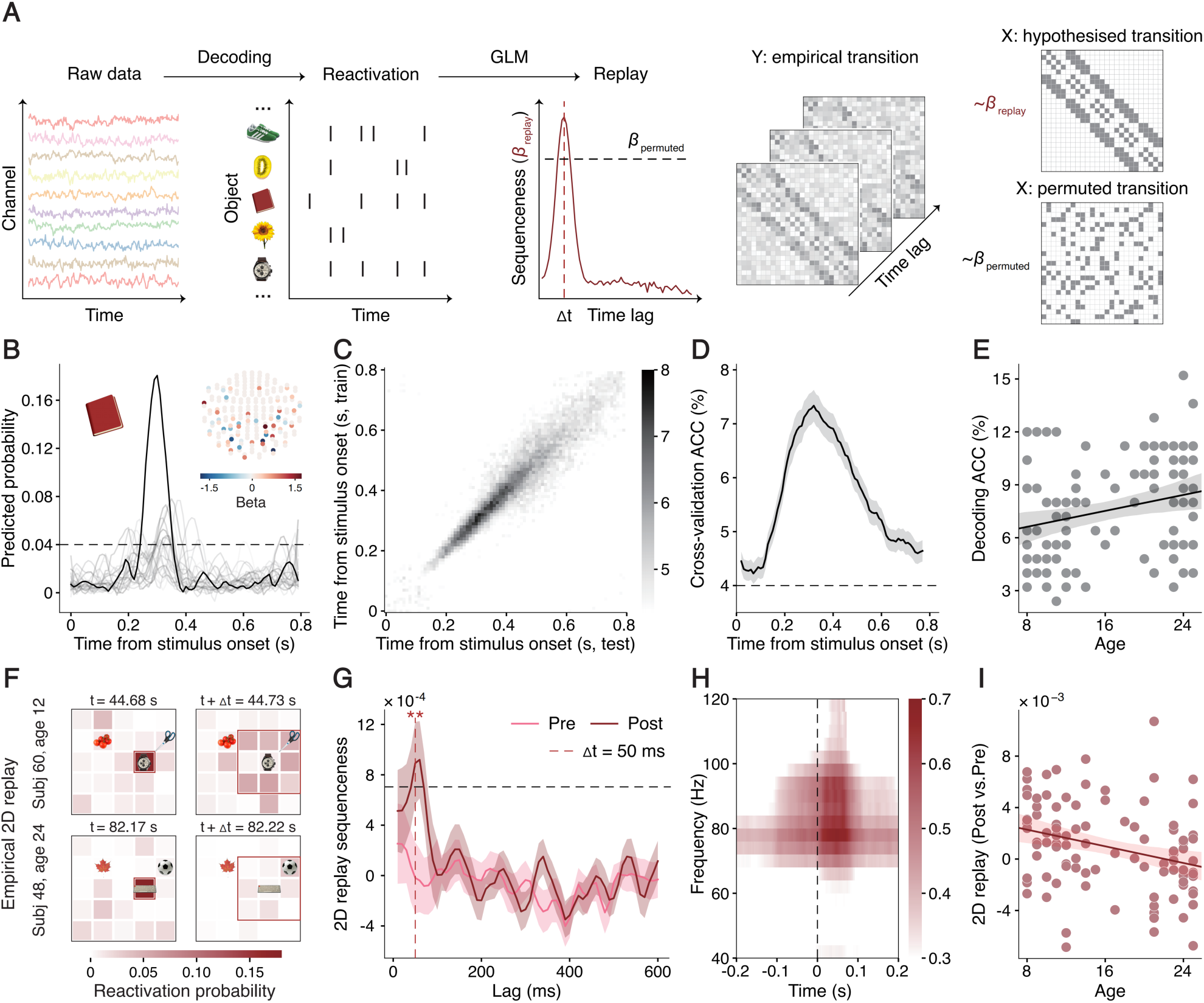
Neural decoding accuracy and task-induced replay across ages. **(A)** Pipeline of neural replay analysis. Neural decoders were trained for each object using task-evoked MEG data. These decoders were then applied to resting-state MEG data, transforming the raw neural signals into time series of reactivation probability, which reflect the likelihood that neural activity corresponds to a specific object at a given time. Replay was quantified by assessing the sequential activation patterns among objects. Specifically, for a given time lag 𝛥𝑡, the reactivation probability of one object was used to predict the future reactivation probability of all other objects. This resulted in empirical transition matrices, representing observed transition probabilities between objects. Replay sequenceness was then defined as the regression coefficient *𝛽_replay_* obtained by comparing these empirical transition matrices with hypothesised transition matrix (**Supplementary Figure 2C)**. Statistical significance was determined by comparing 𝛽*_replay_* from empirical transitions to those from randomly permuted transition matrices (𝛽*_permuted_*). **(B)** Example of neural decoding. Object-specific neural decoders were trained and tested at every 10 ms interval from stimulus onset to 800 ms, using task-evoked MEG data in sensor space. The example illustrates the predicted probabilities for the object “book”, decoded by 25 different decoders trained at 300 ms post-stimulus onset. The probability predicted by the “book” decoder (black line) is higher than those predicted by other decoders. The inset shows the sensor distribution and beta weights for the “book” decoder. **(C)** Decoding accuracy across training and testing time points. Decoding accuracy was highest when training and testing were conducted at the same time point. **(D)** Decoding accuracy over time. Cross-validation results showed that peak decoding accuracy was achieved at 300 ms post-stimulus onset. Error bars represent the group SEM. This peak was used as the basis for selecting neural decoders to detect replay during rest. **(E)** Correlation between decoding accuracy and age. Decoding accuracy (with decoders trained at 300 ms) showed a significant positive correlation with age. The solid line represents the best-fitting line across participants, and the shaded area indicates the 95% confidence interval. Each dot represents an individual participant. **(F)** Example 2D replay sequences. 2D replay is characterised by the sequential reactivation of nearby objects on the hidden 2D map. Representative 2D replay sequences from a younger and an older participant demonstrate that, in both cases, the activation of a starting object at time *t* is followed by adjacent objects after a 50 ms interval (red boxes mark starting and adjacent objects). 2D replay was more pronounced in younger participants compared to older participants. **(G)** 2D replay during rest. 2D replay during post-task rest was significantly higher than expected by chance (black dashed line indicates the permutation threshold) at time lags of 50–70 ms. Additionally, post-task replay was significantly higher than pre-task replay at the 50 ms lag. Error bars represent the group SEM. ‘**’ for *p* < 0.01. See also **Supplementary Figure 2**. **(H)** Extraction of ripple component. A neural component associated with ripple activity was extracted using a generalised eigendecomposition method (Ohki et al., 2024). Epoching this component around replay events revealed increased high-frequency power during replay, indicating replay-related modulation of ripple-band activity. **(I)** Age-related decline in task-induced 2D replay. The change in 2D replay from pre-task to post-task rest (post minus pre) was negatively correlated with age, indicating a decline in task-induced replay in older participants. The solid line represents the best-fitting line across participants, and the shaded area indicates the 95% confidence interval. Each dot represents an individual participant.

Using these decoders, we transformed resting-state MEG data into time series of reactivation probabilities for each object. We defined replay events as sequential patterns of object reactivation, quantified by the “sequenceness” measure from the temporally delayed linear modelling (TDLM) method (Liu et al., 2021). As in Ou et al. (2025), we classified replay experiences as either “1D replay”, reflecting separate consolidation along a single dimension (for instance, “watch → tomato”), or “2D replay”, reflecting adjacency on the hidden 2D map (for instance, “watch → scissors”; see also **Figure 2F**). If both 1D and 2D replays occurred, participants would be consolidating each dimension separately. By contrast, if only 2D replay exists, as similar with Ou et al. (2025), it suggests replay represented a unified 2D map.

We found no evidence of 1D replay in pre-task or post-task rest (see **Supplementary Figure 2**). However, 2D replay did emerge during post-task rest (**Figure 2F–G**). Across all subjects, 2D replay was stronger than chance at 50–70 ms lags, controlling for multiple comparisons (**Figure 2G**). Across both pre-task and post-task rest, 2D replay peaked at a 50 ms lag, and we focused on this lag for all further analyses. Notably, 2D replay was significantly higher during post-task rest than during pre-task rest (paired-sample *t*-test, *t*(97) = 3.065, *p* = 0.003), suggesting that participants organised the inferred information into a 2D map after performing the inference task. Moreover, the increase in 2D replay from pre-task to post-task rest was negatively correlated with age (*r* = −0.300, *p* = 0.003; **Figure 2I**), and this relationship persisted after controlling for decoding accuracy (*r* = −0.293, *p* = 0.004), head movement (*r* = −0.261, *p* = 0.017), head size (*r* = −0.306, *p* = 0.002), memory (*r* = −0.302, *p* = 0.003) and task (*r* = −0.304, *p* = 0.002) performances. Younger participants thus displayed greater reliance on replay to represent the 2D map, whereas older participants engaged in less replay.

Because replay often associates with hippocampal ripples, we also checked whether 2D replay co-occurred with ripple-band (80–120 Hz) power increases. Indeed, 2D replay was accompanied by stronger ripple-band power compared to the global average (paired-sample *t*-test, *t*(97) = 6.189, *p* < 0.001; see also **Supplementary Figure 3**). Using generalised eigendecomposition (Ohki et al., 2024), we extracted a ripple-related component, and time-frequency analysis revealed increased high-frequency power around replay onset (**Figure 2H**). Across participants, this ripple power increase was negatively correlated with age (*r* = −0.242, *p* = 0.016), again indicating heavier replay engagement in younger participants. Taken together, these findings suggest that as individuals mature, they become less dependent on replay during rest for constructing the 2D map, possibly reflecting a shift toward more efficient, stable representations.

### Distinct temporal and spectral characteristics of the DMN remain stable across ages

Beyond replay processes, resting-state cortical dynamics also shape how the brain organises information during rest (Tambini et al., 2010). To investigate how replay–DMN coordination evolves from childhood to adulthood, we first needed to detect activation of resting-state networks (RSNs) from MEG data and examine their intrinsic dynamics. We used the time-delay embedded hidden Markov model (TDE-HMM) method (Vidaurre et al., 2018; see Methods for details). This approach models neural activity as latent states, each defined by a unique covariance matrix describing power and cross-spectra across brain regions. Parameters are estimated through Bayesian inference, yielding both state covariances and activation time-courses.

Specifically, the raw MEG data were reconstructed into source space (**Figure 3A**, see also Methods), parcellated into 38 regions of interest (**Supplementary Table 2**), and the resulting time series were input into TDE-HMM (Gohil et al., 2024). We then estimated power and coherence maps for each inferred state (see also **Supplementary Figure 4A**) and identified known brain networks based on their spatial patterns. To group similar states, we reordered state labels by their transition probabilities (**Supplementary Figure 4B, C**). Ultimately, we identified 12 RSNs. RSN state 1 corresponded to the DMN, primarily involving the medial prefrontal cortex (mPFC) and posterior cingulate cortex (PCC), a pattern robustly replicated across multiple datasets (Van Es et al., 2023). RSN state 2 corresponded to the visual network (VIS), covering the occipital lobe. These two states are highlighted because of their temporal coordination with replay identified in prior literature (Higgins et al., 2021; Ji & Wilson, 2007) and in subsequent analyses. Activation probability time series were generated for each state, and the mutually exclusive nature of these states allowed us to produce a deterministic activation time series.

**Figure 3.**
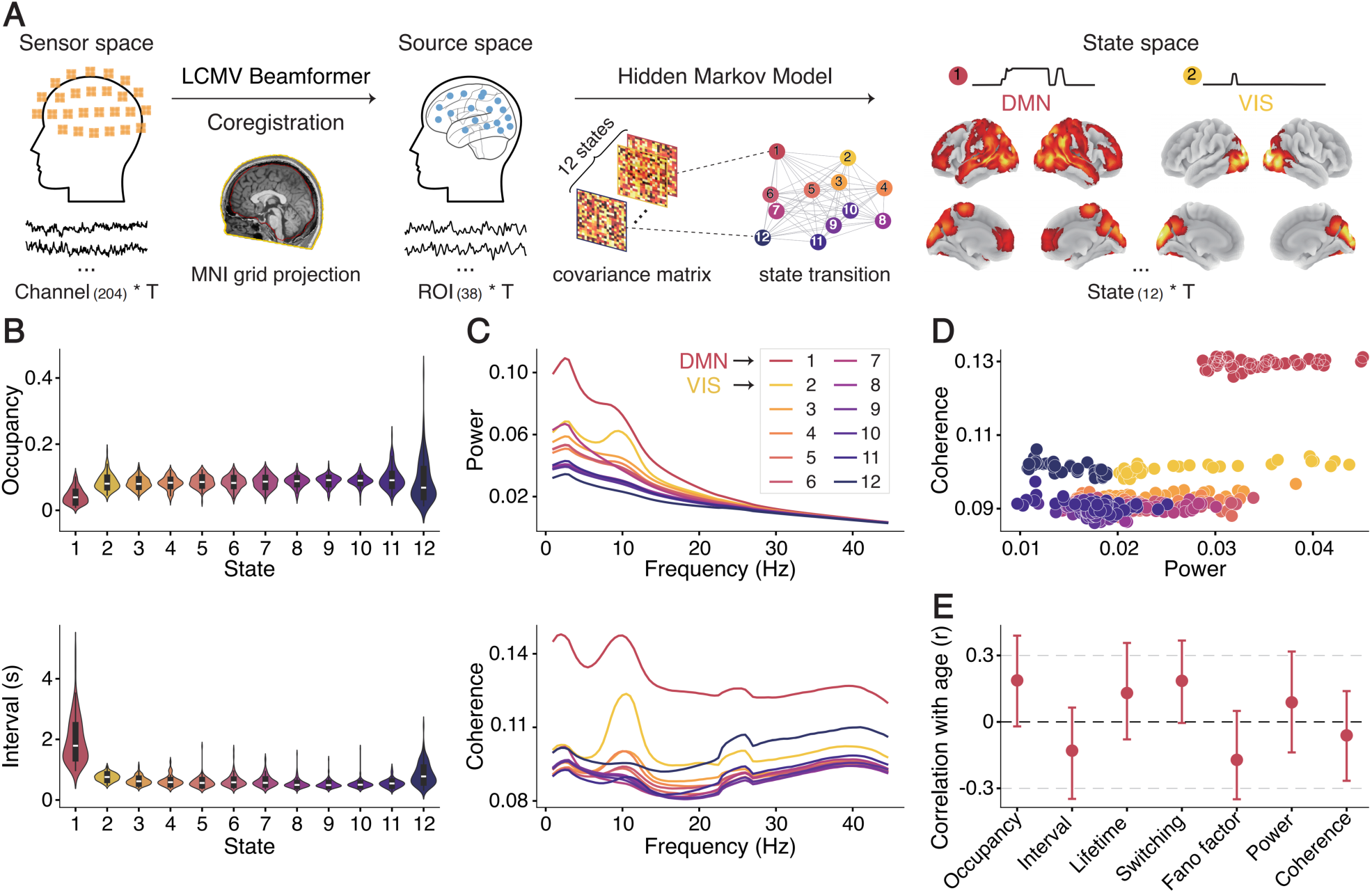
Distinct temporal and spectral characteristics properties of the DMN remain stable across ages. **(A)** Identification of 12 RSNs using the TDE-HMM. Post-task resting-state MEG time series from all planar MEG channels were reconstructed into source space via an LCMV beamformer projection, with individual structural MRI coregistration to improve spatial resolution. The reconstructed data were then parcellated into 38 predefined regions of interest (see **Supplementary Table 2**). The HMM transformed these time series into probabilities of RSN-state activation. The model estimated covariance matrices (characterising the power spectrum within RSN states) and transition probabilities (describing dynamics between RSN states). RSN states were labelled based on their minimised distance. Thresholded power maps for state 1 (left, red, DMN) and state 2 (right, yellow, VIS) are presented. See also **Supplementary Figure 4A.** **(B)** Temporal characteristics of RSN-state activation. RSN states are mutually exclusive, with one state having the highest activation probability at each time point, which can be used to generate binarised state activation time series. This allows fractional occupancy and mean interval for each RSN state to be quantified. The DMN exhibited the lowest occupancy, and the longest intervals compared to other RSN states, with all paired-sample *t*-tests *p* < 0.001 after family-wise error correction. Violin plots display these distributions, with internal boxplots marking medians and quantiles across participants. **(C)** Spectral characteristics of RSN-state activation. The DMN demonstrated the highest power and coherence across frequencies, featuring peaks in the delta/theta and alpha bands. **(D)** Relationship between RSN power and coherence. Scatterplot showing the wideband, subject-averaged power and coherence per state. Each dot represents a value from specific brain regions in source-space data. The DMN maintained high power and coherence concurrently, distinguishing it from other RSN states. **(E)** No significant relationship between general DMN characteristics and age. Key temporal (fractional occupancy, mean interval) and spectral (power, coherence) characteristics of the DMN showed no significant correlation with age. Dots represent observed correlation values across participants, with error bars indicating 95% confidence intervals. See **Supplementary Table 3** for correlation statistics.

We then summarised the temporal and spectral characteristics of these RSNs. Temporally, we used the state activation time series to calculate fractional occupancy (the proportion of time spent in each state) and mean interval time (the average duration between consecutive state visits). Among all states, the DMN showed the lowest fractional occupancy and the longest mean interval (**Figure 3B**). Spectrally, for each state, we computed mean power and mean coherence across all brain regions and frequencies. The DMN exhibited the highest overall power and coherence, with peaks in the delta/theta and alpha bands (**Figure 3C, D**), and these patterns were replicated in factorised frequency bands (see **Supplementary Figure 4D**). Together, these observations highlight the distinctiveness of DMN relative to other RSNs across all subjects, including children, consistent with previous findings in adults (Higgins et al., 2021).

Notably, none of these general DMN properties correlated with age (**Figure 3E**). The temporal measures, including fractional occupancy, mean interval time, mean lifetime (average duration of state activation), switching rate (average number of state activations), and Fano factor (temporal irregularity), showed no significant relationships with age (all *p* > 0.1). Likewise, the spectral measures, including power (*r* = 0.088, *p* = 0.387) and coherence (*r* = −0.061, *p* = 0.551), also did not correlate with age. This suggests that the fundamental temporal and spectral characteristics of the DMN remain stable across development.

### Replay–DMN alignment grows with age to support schema-based knowledge

We next asked whether replay interacts with RSN activity, particularly the DMN, and how such replay–DMN coordination evolves through development. We also examined whether this alignment underpins grid-like schema representations in the EC, testing our central question of how replay transitions from direct associations to aligning new experiences with existing schemas.

Consistent with studies in adults (Higgins et al., 2021), we found that replay events were temporally aligned with increases in DMN activation across all participants (see **Figure 4A** for a representative example). At the group level, we observed a transient increase in activation probability in both the DMN and the VIS around replay events, peaking approximately 30 ms after replay onset (**Figure 4B**). Sign-flipping permutation tests confirmed the significance of these increases (*p* < 0.001; see Methods for details).

**Figure 4.**
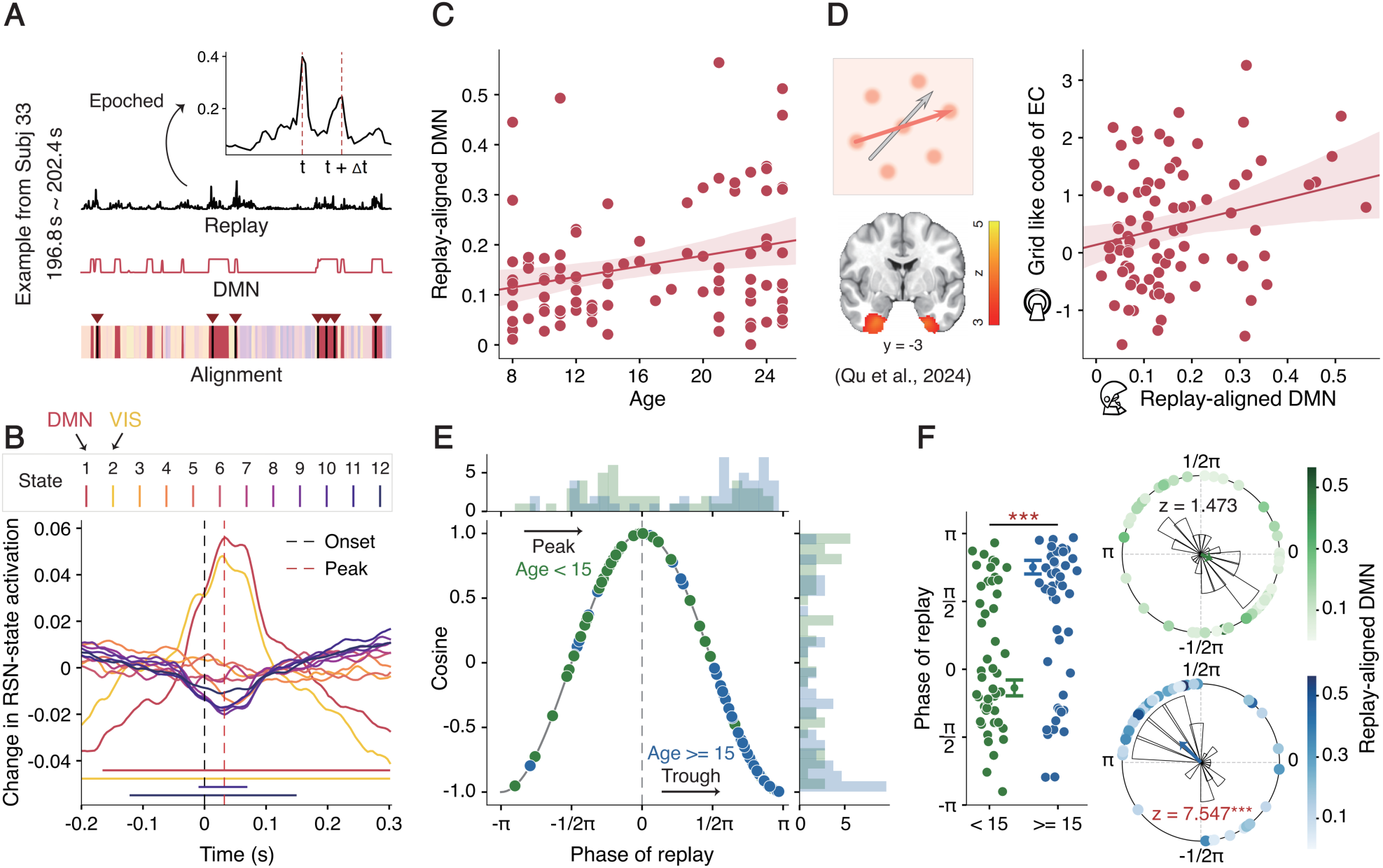
Replay–DMN alignment becomes stronger and more synchronised with age. **(A)** Example of replay–DMN alignment. An example from Subject 33 (196.8 s to 202.4 s) illustrates the temporal coupling between replay and DMN activation. Replay quantifies the extent to which strong reactivation of one object is immediately followed by strong reactivation of subsequent objects. The black line represents the summed reactivation probability across all possible object transitions, with peaks marking replay onsets. The inset shows reactivation probability after epoching around replay onsets, revealing a peak at time 𝑡, followed by a secondary increase at 𝑡 + 𝛥𝑡. The red line represents DMN activation. Their alignment demonstrates how replay events (red inverted triangles) were consistently accompanied by DMN activation (red colour). **(B)** Changes in RSN-state activation around replay events. Replay events accompanied by a transient increase in RSN-state activation, particularly in the DMN and VIS. This increase was observed from −0.2 s to 0.3 s relative to replay onset (dashed black line), peaking around 30 ms after replay onset (dashed red line). Significance bars indicate time windows where RSN-state activation was significantly modulated by replay events (*p* < 0.001, permutation tests). **(C)** Replay-aligned DMN activation positively correlates with age. The solid line represents the best-fitting line across participants, and the shaded area indicates the 95% confidence interval. Each dot represents an individual participant. **(D)** Replay-aligned DMN activation positively correlates with grid-like code in the EC. Grid like code in EC was found in Qu et al. (2025) with the same set of participants using fMRI (see Methods for details). The solid line represents the best-fitting line across participants, and the shaded area indicates the 95% confidence interval. Each dot represents an individual participant. **(E)** Theta phase coupling of replay events varies with age. Replay events became more phase-locked to the trough of theta oscillations with age. Participants were split into two groups based on median age for visualisation. In younger participants (age < 15, green), replay events were distributed across all phases, whereas in older participants (age ≥ 15, blue), replay events were preferentially clustered in the theta trough. Each dot represents the mean theta phase of replay onsets for a participant. **(F)** Stronger replay-aligned DMN activation occurs at the theta trough. Replay events showed a significant difference in phase distribution between younger and older participants (left panel, Watson-Williams test). Error bars represent the group SEM. Further analysis revealed that in younger participants, replay events were not systematically locked to specific theta phases (right panel, Rayleigh test). In contrast, in older participants, there was a significant clustering of replay events in the theta trough. More importantly, stronger replay-aligned DMN activation was observed when replay events occurred in the theta trough. Each dot represents the mean theta phase of replay onsets for a participant; dot colour intensity reflects the strength of replay-aligned DMN activation (darker colours indicate stronger activation). ‘***’ for *p* < 0.001.

Next, we investigated whether replay-aligned DMN activation correlates with age. We defined replay-aligned DMN activation as the baseline-corrected DMN activation probability, centred on replay onsets (see Methods for details). For each participant, we extracted replay-aligned DMN activation at the group-level peak. We observed a significant positive correlation between age and replay-aligned DMN activation (*r* = 0.281, *p* = 0.005; **Figure 4C**), which persisted after controlling for head movement (*r* = 0.316, *p* = 0.004), head size (*r* = 0.287, *p* = 0.004), memory (*r* = 0.277, *p* = 0.006) and task (*r* = 0.253, *p* = 0.012) performances. Thus, as individuals mature, the coupling between replay events and DMN activity becomes stronger. Notably, this relationship was absent in other RSNs, including the VIS (see **Supplementary Figure 5A** for control analysis of the VIS), and was significantly stronger for the DMN than for the VIS, as confirmed by Steiger’s *Z*-test (*Z* = 2.546, *p* = 0.011).

This strengthened replay–DMN coordination appears to facilitate aligning newly replayed content with pre-existing schemas. To support this idea, we examined a subset of participants (N = 85) who completed both the MEG task and a subsequent fMRI-based map inference task (Qu et al., 2025). In those subjects, we identified grid-like codes in the EC using fMRI, reflecting a 2D schematic representation of the task space. The strength of the EC grid-like code was significantly above chance (one-sample *t*-test, *t*(84) = 4.197, *p* < 0.001) and positively correlated with age (*r* = 0.226, *p* = 0.038), confirming earlier results (Qu et al., 2025). Crucially, in the same participants, replay-aligned DMN activation predicted the strength of the EC grid-like code (*r* = 0.243, *p* = 0.025; **Figure 4D**). An exploratory mediation analysis indicated that replay-aligned DMN activation during rest may mediate the relationship between age and grid-like code strength (indirect effect, *p* = 0.069).

To probe the neural mechanism underpinning this developmental shift, we explored how replay becomes increasingly coordinated with large-scale cortical dynamics. One potential mechanism is phase-based timing in the theta band, which has been linked to hippocampal–cortical communication and memory integration (Fujisawa & Buzsáki, 2011; Heusser et al., 2016; Veselic et al., 2025). We hypothesised that the stronger replay–DMN alignment seen in older participants reflects more precise timing of replay relative to the ongoing theta rhythms, allowing more effective embedding of replayed content. We therefore analysed the theta phase (2–6 Hz, derived by non-negative matrix factorisation, Higgins et al., 2021; see also **Supplementary Figure 4D**) at which replay events occurred for each participant. Among all source-space regions, the PCC, a core DMN node, showed the highest theta power, so we focused on that region.

To address sign ambiguity in source-space MEG data (Vidaurre et al., 2016), we applied a sign-flipping procedure based on the power of theta troughs and peaks (see Methods for details) and validated it using a replay event-related phase-amplitude coupling analysis (see **Supplementary Figure 6**). We found a significant correlation between age and the replay event phase (circular-linear correlation, *ρ* = 0.300, *p* = 0.012), demonstrating that older participants had replay events more aligned with the theta trough.

We further characterised this developmental transition by splitting participants at the median age. Younger participants (age < 15) had replay onsets near the theta peak, whereas older participants (age ≥ 15) were more aligned with the theta trough (**Figure 4E**). A Watson-Williams test confirmed that mean replay phases differed significantly between groups (*F*(1, 96) = 36.013, *p* < 0.001; **Figure 4F**, left), and a Rayleigh test showed that only older participants exhibited significant phase locking of replay events to the theta trough (*z* = 7.547, *p* < 0.001). Younger participants displayed no significant phase locking (*z* = 1.473, *p* = 0.230; **Figure 4F**, right).

To confirm that global theta oscillations alone did not drive the observed phase alignment, we conducted a control analysis by randomly shuffling decoder weights (see Methods for details). This generated random reactivations unrelated to actual replay, allowing us to test whether such random events would align with the theta trough. We found no correlation between age and replay phases derived from shuffled decoders (circular-linear correlation, *ρ* = 0.159, *p* = 0.290), indicating that the observed phase alignment is specific to genuine replay events rather than general theta dynamics.

Importantly, in older participants, replay events closer to the theta trough were accompanied by stronger DMN activation (circular-linear correlation, *ρ* = 0.395, *p* = 0.022; **Figure 4F**, right), suggesting a functional role for theta phase in coordinating replay–DMN interactions. This effect did not appear in younger participants (*ρ* = 0.198, *p* = 0.384).

Overall, these findings reveal that replay–DMN alignment becomes increasingly synchronised with age, with theta-phase locking emerging as a key mechanism. As subjects mature, replay events are timed more precisely to the theta trough, leading to stronger DMN engagement and potentially more effective embedding of new information into established grid-like schema representation.

### Age-related strengthening of DMN connectivity reduces replay reliance and improves inference

Building on our observation that replay–DMN alignment becomes more pronounced and precisely timed with age, we examined whether broader connectivity features of the DMN contribute to these developmental changes. Specifically, we tested whether resting-state functional connectivity (rsFC) within the DMN, measured via fMRI, might support neural replay (observed in MEG) and inference performance.

Although the general temporal and spectral properties of the DMN from MEG-based analyses were stable across ages, earlier work has reported that rsFC within the DMN increases with development (Fair et al., 2009; Fan et al., 2021), potentially reflecting adaptive changes that facilitate memory processes (Sestieri et al., 2011; Xia et al., 2022). It is possible that age-related strengthening of DMN rsFC leads to more efficient representations of the cognitive map and thereby decreases reliance on replay mechanisms.

To test this idea, we employed connectome-based predictive modelling (CPM) (Scheinost et al., 2019; Shen et al., 2017) to determine whether rsFC within the DMN, derived from resting-state fMRI, could predict changes in 2D replay measured by MEG (**Figure 5A**; see Methods for details). Specifically, we constructed individual rsFC matrices by correlating the average BOLD signals from each voxel in the DMN, as defined by the Schaefer 400 atlas (Schaefer et al., 2018). We then performed leave-one-out cross-validation (LOOCV) at the participant level, applying feature selection within each training set to build a linear model predicting 2D replay strength in the left-out participant.

**Figure 5.**
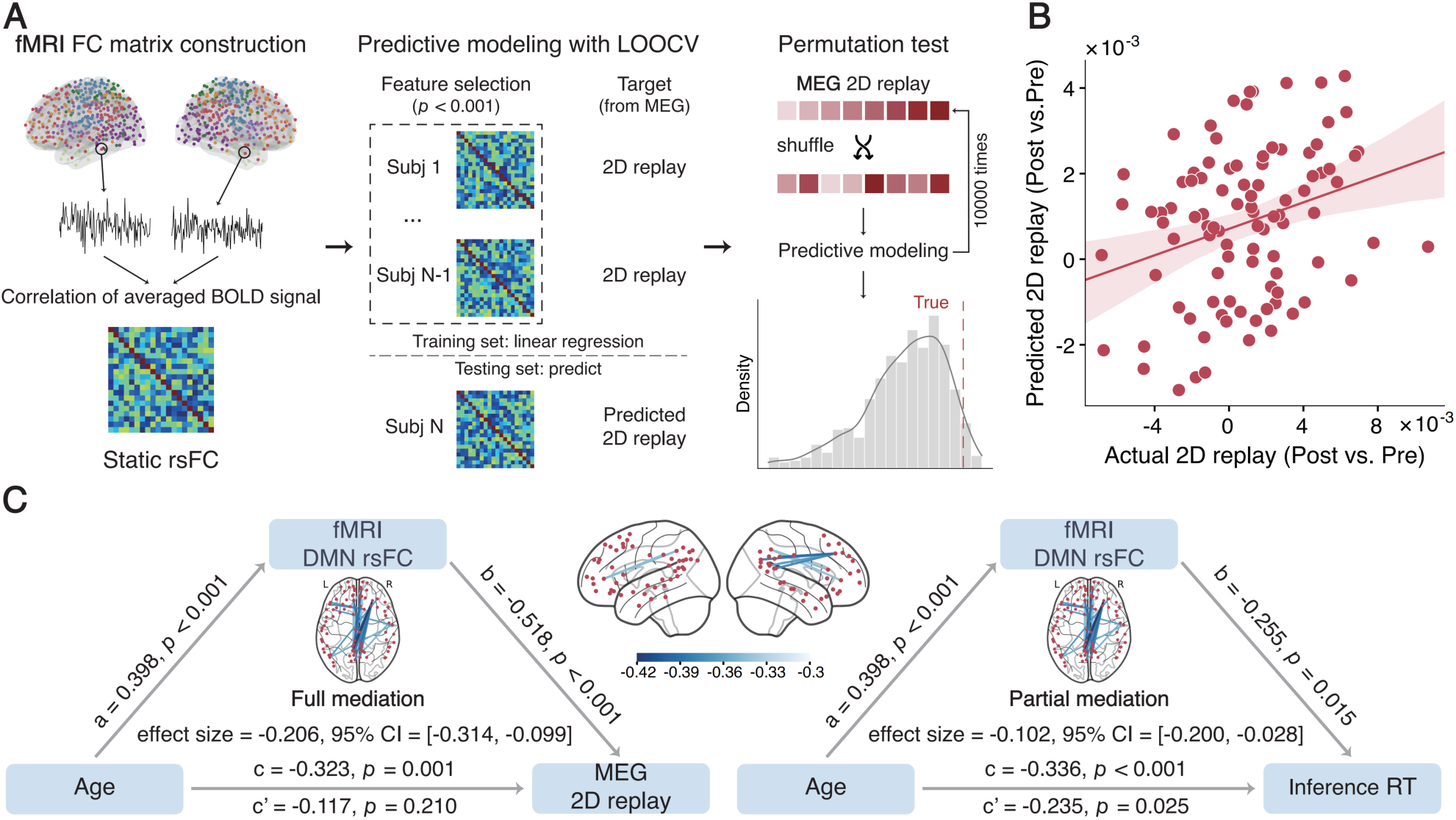
Age-related strengthening of DMN connectivity reduces replay reliance and improves inference. **(A)** Connectome-based predictive modelling of rsFC for predicting task-induced 2D replay. Resting-state fMRI data were used to construct rsFC matrices by correlating averaged BOLD signals from each voxel within the DMN, defined by the Schaefer 400 atlas. LOOCV was performed at the participant level, with feature selection conducted within the training set (𝑁 − 1 participants). A linear regression model was then built to predict the 2D replay strength of the left-out participant. Model performance was evaluated by the Spearman correlation between predicted and observed replay values, with permutation tests (10,000 iterations) used to assess statistical significance. **(B)** DMN rsFC predicts MEG-based task-induced 2D replay. The predicted replay strength positively correlated with the actual replay. The solid line represents the best-fitting line across participants, and the shaded area indicates the 95% confidence interval. Each dot represents one validation data point. Permutation tests further validated the significance. **(C)** Mediating analyses of the DMN rsFC in fMRI. The strength of DMN rsFC fully mediated the age-related decrease in 2D replay (left panel) and partially mediated the age-related improvement in inference efficiency, reflected by faster reaction times (right panel). Insets show the spatial distribution of robust features (selected with frequency above 0.9) contributing to the predictive model.

Our results showed that DMN rsFC significantly predicted the task-induced replay (Spearman’s *ρ* = 0.275, *p* = 0.007; **Figure 5B**). Permutation tests supported the statistical significance of this prediction (*p* = 0.034). In contrast, rsFC within the VIS did not predict replay sequenceness (Spearman’s *ρ* = −0.031, *p* = 0.766, see **Supplementary Figure 5B**), and the difference between DMN and VIS predictions was statistically significant (Steiger’s *Z* = 2.221, *p* = 0.026). Further analysis of whole-brain rsFC revealed that connectivity within and across the DMN contributed most strongly to the predictive model (see **Supplementary Figure 5C**). Feature selection in CPM revealed specific DMN rsFC connections that robustly predicted 2D replay; averaging these connections yielded a summary index of DMN rsFC, which showed a significant positive correlation with age (*r* = 0.313, *p* = 0.002). This finding is consistent with earlier reports that DMN connectivity strengthens with development.

Finally, we conducted mediation analyses to determine whether DMN rsFC could explain the relationship between age, task-induced replay, and inference efficiency. We found that DMN rsFC fully mediated the age-related reduction in 2D replay (indirect effect size, *d* = −0.206, 95% CI [−0.314, −0.099], *p* < 0.001; **Figure 5C**, left panel). Thus, as DMN connectivity increases with age, it reduces reliance on replay for representing the 2D map. DMN rsFC also partially mediated the age-related improvement in inference efficiency, reflected in faster reaction times (indirect effect size, *d* = −0.102, 95% CI [−0.200, −0.028], *p* = 0.033; **Figure 5C**, right panel). This suggests that greater DMN connectivity promotes more efficient inferential behaviour.

To rule out potential age-related confounds such as task engagement, fatigue, head motion, head size, or processing speed, we performed thorough complementary analyses, none of which explained the observed relationships (see **Supplementary Note 1** and **Supplementary Figures 7**–10 for details). In summary, by integrating MEG and fMRI data, our results show that age-related increases in DMN connectivity, together with more precise replay–DMN alignment, are crucial for reducing reliance on replay alone and enhancing inference abilities.

## Discussion

Our findings provide novel insights into how neural replay and its coordination with large-scale networks change over development. In our sample, younger participants showed a stronger reliance on replay during post-task rest, whereas older participants demonstrated increasingly precise replay–DMN coordination, which in turn predicted the strength of an entorhinal grid-like code. This pattern indicates a gradual shift from replay occurring on its own to replay events timed to specific moments in distributed cortical activity (i.e. coupled to ongoing network oscillations) as individuals mature.

Broadly, there are two conceptual modes for representing relationships among experiences (Behrens et al., 2018). One involves building a direct associative map of inter-item links—for example, a successor representation constructed by tracking sequential experiences (Barron et al., 2020; Stachenfeld et al., 2017). The other entails incorporating new information into an existing schema, allowing rapid assimilation of knowledge into an abstract framework (Tse et al., 2007; Van Buuren et al., 2014). Emerging evidence suggests that hippocampal replay can support both mapping strategies: it can reinforce local associative links and also help integrate memories into schema-like structures depending on context (Bakermans et al., 2025; Evans & Burgess, 2019; Ou et al., 2025). Consistent with this, children who have not yet formed coherent grid-like schemas tend to rely more on hippocampal mechanisms for inference (Qu et al., 2025). In line with previous observations, our results show that younger participants used replay to integrate experiences into a 2D cognitive map—effectively “stitching together” the learned pairwise associations—perhaps to stabilise inferred relationships at an age when schematic representations were still rudimentary. Notably, this age difference in replay usage cannot be explained by differences in neural decoding fidelity, which in fact improved with age (presumably due to more refined sensory representations in older individuals) (Fandakova et al., 2019). Thus, replay appears to compensate for underdeveloped schema frameworks in younger individuals, acting as a mechanism to solidify new knowledge when abstract cognitive maps are not yet fully formed.

A key finding of our study is that although replay events became less frequent in older participants, they were more tightly aligned with concurrent DMN activity. This enhanced replay–DMN synchrony was strongly associated with the emergence of grid-like (schema-related) representations in the EC, suggesting that timing replay to coincide with high-excitability phases of DMN activity helps incorporate reactivated experiences into broader knowledge networks (Kaefer et al., 2022; Masís-Obando et al., 2022). Physiologically, older participants showed more precise timing of replay events at the trough of the DMN theta rhythm localised in the PCC, a hub region with dense connectivity to the hippocampus and medial temporal lobe (Andrews-Hanna et al., 2010; Greicius et al., 2009). The PCC is well known to integrate memory-related signals and coordinate top–down influences across cortical areas (Alexander et al., 2023; Goddard et al., 2022). Recent studies in rodents have further revealed that retrosplenial cortex (RSC) activity, which shares functional properties with PCC in humans, becomes time-locked to hippocampal ripples during sleep (Pedrosa et al., 2022; Swanson et al., 2025), indicating that hippocampal–cortical interactions are modulated by ongoing cortical oscillations. Consistent with this idea, the trough of the theta cycle is associated with heightened cortical excitability for hippocampal inputs (Buzsáki & Draguhn, 2004; Fuentemilla et al., 2010; Sirota et al., 2008), potentially enabling more effective embedding of replayed content into distributed networks. Notably, human theta-band rhythms are generally slower than those in rodents. In this study, we defined the cortical theta range as 2–6 Hz based on data-driven spectral decomposition, and we found that replay events in older participants aligned with the DMN’s theta-phase dynamics rather than with hippocampal intrinsic theta oscillations (Kragel et al., 2020; Solomon et al., 2019). This implies that maturation brings about a more network-level form of memory integration (Kaefer et al., 2022), wherein hippocampal replay is coordinated with global cortical rhythms (as opposed to being locked only to local hippocampal theta). Future work should investigate how this precise coupling of replay to cortical theta emerges at the circuit level during development (Fernández-Ruiz et al., 2017; Xiao et al., 2024).

In addition to changes in oscillatory coordination, we observed that resting-state functional connectivity within the DMN increased with age, and this change could explain the decreasing dependence on replay and the improvement in inference performance across development. Although the temporal signatures of DMN activation (as measured with MEG) remained stable with age, the fMRI-based connectivity measures indicated that the DMN became more cohesive and strongly interconnected in older participants. A more strongly connected DMN may provide a robust scaffold for integrating reactivated memory content into existing knowledge structures (Xiao et al., 2025). Taken together, our results highlight an interplay between replay, the maturation of cortical networks, and the timing of neural events that collectively supports the development of schema-based inference abilities.

These findings have broader implications for learning and memory in childhood and adolescence. The heavier reliance on replay in younger individuals to stabilise new information suggests that memory replay-based strategies—such as active recall or spaced repetition (Antony et al., 2017; Karpicke & Roediger, 2008; Wiklund-Hörnqvist et al., 2021)—might be particularly beneficial for children and pre-adolescents. In contrast, older adolescents and young adults, who exhibit stronger DMN connectivity and more precise replay–DMN timing, may benefit more from schema-focused learning approaches that leverage existing knowledge frameworks and abstract reasoning (Aghayan Golkashani et al., 2021; Ramey et al., 2024; Van Kesteren et al., 2012). Further investigation of individual differences in replay dynamics and DMN network maturity could help tailor educational strategies to different developmental stages.

In conclusion, our study delineates a developmental trajectory in which an early reliance on replay gradually transforms to integrate with DMN mechanism characterised by precise theta-phase coupling and enhanced DMN connectivity. This shift appears to underlie the formation of entorhinal grid-like schema representations and supports more efficient inferential reasoning in mature cognition. By elucidating how replay processes become coordinated with brain-wide networks as children age, we provide a mechanistic framework for understanding cognitive development and open new avenues for interventions to optimise learning and memory consolidation across development.

## Methods

### Participants

The study comprised multiple sessions, including off-scan behavioural training, MEG scanning, and MRI scanning. A total of 106 right-handed participants aged 8–25 years (mean ± SEM: 16.45 ± 0.61 years; 51 females, see also **Supplementary Table 1**) were recruited after qualifying in the training session. All participants had normal or corrected-to-normal vision and reported no neurological disorders, brain injuries, or developmental disabilities.

After the MEG scanning session, data from 98 participants were retained for analysis; eight were excluded due to excessive head movement. The final sample for MEG analyses had similar demographics (mean ± SEM: 16.40 ± 0.64 years; 49 females). Behavioural data and resting-state MEG data from these 98 participants were used for all subsequent analyses. Of these, 96 participants completed the resting-state fMRI scanning session. Following the resting-state fMRI scanning session, 85 participants who completed the MEG experiment also took part in a task fMRI study probing the grid-like code of the 2D knowledge map through development, as detailed in Qu et al. (2025).

Prior to participation, informed consent was obtained from all participants or their guardians, along with two scanning safety checklists. The study adhered to the Declaration of Helsinki and was approved by the Institutional Review Board of the Faculty of Psychology at Beijing Normal University (ethics number: IRB_A_0051_2021002). Participants received monetary compensation upon completing the experiment. Children were encouraged to participate through laboratory visits and developmental guidance.

### Experimental design

The experiment consisted of three main sessions: off-scan behavioural training, MEG scanning, and MRI scanning.

In the off-scan training session, participants underwent extensive behavioural training to learn a cognitive map. Twenty-five objects were arranged in an unseen 5 × 5 2D grid, representing two attributes: attack power (AP) and defence power (DP). Participants learned the power levels of each object along these two dimensions separately through pairwise comparisons. For example, in a learning trial for the attack power dimension, two objects (e.g., a key and a pepper) were presented simultaneously, and participants chose the object with higher power. They gradually learned the true sequence through trial-and-error iterations. Importantly, the paired objects differed by one power level, resulting in 100 unique pairs for each dimension.

From the participants’ perspective, the learned relationships could be represented as multiple one-dimensional (1D) sequences for each dimension or as a single integrated 2D map combining both dimensions. The learning order of the two dimensions and key-press assignments was balanced between participants. Extensive training over multiple days ensured that participants reached at least 80% accuracy, qualifying them for the subsequent scanning sessions. The overall training performance is summarised in **Supplementary Note 1**, while detailed descriptions of the training procedure can be found in Qu et al. (2025).

The MEG scanning session involved a five-minute pre-task rest, a map-based inference task, and a five-minute post-task rest. During the map-based inference task, participants were asked to infer which of two randomly selected objects had higher power in either the attack or defence dimension. Although this inference could be made using either 1D or 2D representations, utilising the integrated 2D map was expected to be more efficient. The task required participants to compare objects that were not directly adjacent in the learned sequences, encouraging the utilisation of the 2D map structure.

In each trial (**Figure 1B**), two objects were presented sequentially on the screen for 1.5 s each, with an inter-stimulus interval of 0.4–0.6 s. After both objects were shown, a decision cue indicated the dimension (attack or defence) for comparison. Participants had up to 2 s to respond by choosing the object with higher power on the cued dimension. An inter-trial interval of 0.4–0.6 s followed each response. The task consisted of 240 trials, divided into eight blocks (four blocks for each dimension). The inference order of the two dimensions and key-press assignments was balanced between participants, and object locations were counterbalanced within participants.

The MRI scanning session included T1-weighted structural imaging and a five-minute resting-state fMRI scan. Participants were instructed to keep their eyes open during all resting-state scans. Following these sessions, most participants proceeded to task-based fMRI scanning, which included additional cognitive map tasks detailed in Qu et al. (2025). These tasks involved more complex inferences requiring integration across dimensions and learning relationships with newly introduced objects.

Throughout the experiment, tasks were framed within a “monster tournament” storyline to engage participants and facilitate understanding. Clear instructions were provided, and participants confirmed their comprehension before entering the scanner. Post-scan debriefings ensured that all participants, including the youngest, had understood and completed the tasks as intended. The MEG and resting-state fMRI sessions lasted approximately one hour per participant. Also, care was taken to ensure that all participants, including the youngest, fully understood the tasks, minimized fatigue during scanning, and controlled for other age-related confounding factors (see also **Supplementary Note 1**).

### MEG Data acquisition and preprocessing

MEG data were collected using an Elekta Neuromag TRIUX system (Elekta, Helsinki, Finland), which comprises a sensor array of 102 magnetometers and 204 planar gradiometers. Data were sampled at a rate of 1000 Hz. Participants were seated comfortably in a magnetically shielded room to minimise environmental noise and artefacts. Head position was continuously monitored using head-position indicator coils, allowing for real-time tracking relative to the sensor array. The MEG scanning session began and ended with five-minute resting-state periods, interleaved with eight task runs, each lasting approximately three minutes.

MEG data preprocessing followed established protocols (Higgins et al., 2021; Liu et al., 2021), with adjustments made to accommodate the computational requirements of different analyses. Initial preprocessing involved the use of MaxFilter software (Elekta) to correct for head movement artefacts and reduce external noise through signal-space separation. A 0.5 Hz high-pass filter was applied to remove slow drifts, and notch filters were used to eliminate power line interference. The data were then down-sampled to 400 Hz.

Automated algorithms were employed to identify bad segments and channels. Independent component analysis (ICA) was performed separately on concatenated task and concatenated rest data to identify and remove artefactual components based on spatial topography, frequency spectra, and temporal patterns. After artefact removal, the data were further processed according to the specific requirements of subsequent analyses.

For replay detection, the preprocessed data were down-sampled to 100 Hz to reduce computational load while preserving temporal resolution sufficient for replay analysis.

For resting-state network modelling using the HMM, the data were coregistered with individual structural MRI scans to enable precise source localisation. The coregistered data were down-sampled to 250 Hz and bandpass-filtered between 1–45 Hz to focus on frequencies relevant to resting-state networks (Gohil et al., 2024). Source reconstruction was performed using a linearly constrained minimum variance (LCMV) beamformer, and the reconstructed signals were parcellated (taking the first principal component) into 38 predefined regions of interest (ROIs) based on the Harvard-Oxford atlas (Colclough et al., 2016; see **Supplementary Table 2**). To correct for source leakage, symmetric orthogonalisation was applied (Colclough et al., 2015). Dipole sign ambiguity in the source-reconstructed data was resolved using a greedy algorithm that optimised lagged partial correlations across time lags (Vidaurre et al., 2016), ensuring consistent signal alignment across participants.

### MRI Data Acquisition and Preprocessing

MRI data were acquired using a Siemens Prisma 3 T MRI scanner with a 32-channel head coil. T1-weighted structural images were collected using a 3D magnetisation-prepared rapid acquisition gradient echo sequence with the following parameters: repetition time (TR) = 2.53 s, echo time (TE) = 2.98 ms, flip angle = 7°, field of view (FoV) = 256 mm, slice thickness = 1 mm, and voxel size = 0.5 × 0.5 × 1 mm³. Functional images were acquired using a gradient-echo echo-planar imaging sequence (Ogawa et al., 1990) with TR = 3 s, TE = 30 ms, flip angle = 90°, FoV = 192 mm, slice thickness = 3 mm, and voxel size = 3 × 3 × 3 mm³. The five-minute resting-state scan captured 100 whole-brain volumes. We acquired images with a 30° tilt relative to the anterior-posterior commissure line to minimize signal loss in the orbitofrontal cortex (Weiskopf et al., 2006).

MRI data were preprocessed using fMRIPrep (version 20.2.7; Esteban et al., 2019), ensuring standardised and reproducible results. Preprocessing steps included skull-stripping, bias-field correction, and spatial normalisation to the MNI152NLin2009cAsym template at 2 mm resolution for the structural images. Functional data preprocessing involved motion correction, slice-timing correction, and coregistration with T1-weighted images. Spatial distortion correction was performed using the SyN-based susceptibility distortion correction method. Confound regressors, including head motion estimates and signals from cerebrospinal fluid and white matter, were calculated for later denoising. Outputs were generated in standard MNI space for group-level analyses.

### Replay detection

Replay detection was conducted using a temporally delayed linear modelling (TDLM) approach adapted from Liu et al. (2021). The procedure involved two main steps: neural decoding to identify object-specific neural patterns during the task, and detection of replay sequences during rest based on these patterns (**Figure 2A**).

In the neural decoding step, object-specific decoders were trained using task MEG data, as no independent functional localiser was implemented. The task comprised 240 trials, with each object presented approximately 19 times. Sparse logistic regression classifiers with L1 regularisation were employed to identify neural patterns corresponding to each object. Data from all MEG planar channels were epoched from stimulus onset to 800 ms and used for decoding, with 10 ms time bin.

To determine the optimal time point for classifier training, a five-fold cross-validation was conducted, assessing classification accuracy at every 10 ms interval. Classifiers trained and tested at the same time point (i.e., along the diagonal of the temporal generalisation matrix) showed peak accuracy at 300 ms after stimulus onset (**Figure 2D**). Consequently, object-specific classifiers were retrained on the full task dataset at this time point for subsequent replay detection.

The trained classifiers were then applied to the resting-state MEG data to obtain object reactivation probability time series. Replay was defined as the sequential reactivation of specific objects in patterns reflecting the learned relationships. Two types of replay were investigated: two-dimensional (2D) replay and one-dimensional (1D) replay. In 2D replay, sequential reactivation occurred among objects adjacent on the 2D map, capturing the integrated representation of both dimensions (**Supplementary Figure 2A**). In contrast, 1D replay involved sequential reactivation along a single dimension (attack or defence), excluding neighbours on the 2D map to distinguish from 2D replay.

Replay quantification involved applying general linear regression to model the reactivation probabilities of objects at time 𝑡 + 𝛥𝑡, based on the reactivation at time 𝑡, across all time points, with equation (1).

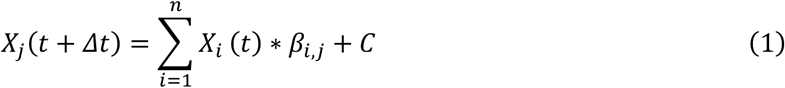

𝑋*_j_*(𝑡 + 𝛥𝑡) is the reactivation probability of object 𝑗, at time 𝑡_0_ + 𝛥𝑡, 𝑋*_i_*(𝑡) is the reactivation probability of object 𝑖, at time 𝑡. 𝛽*_i,j_* represents the empirical transition matrix capturing observed transitions (with shape 𝑁*_object_* × 𝑁*_object_*), and 𝐶 is a constant term.

The empirical transition matrix was predicted from the hypothesised transition matrices while controlling for confounds such as autocorrelation and mean activity. Each hypothesised transition matrix represented every possible transition permitted in either 2D or 1D replay and used binary entries to mark transferable transitions (set to one) and non-transferable transitions (set to zero). For instance, in **Supplementary Figure 2C** (left), the transition “watch → kiwifruit” is given a value of one, whereas “watch → basketball” is given a value of zero in the hypothesised 2D replay transition matrix. The complete equation for modelling the empirical matrix is provided below.

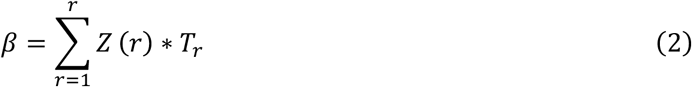

where *𝑇_r_* represented hypothesised transition matrices, and 𝑍(𝑟) was beta coefficient, representing the strength of alignment between empirical and hypothesised transitions, were interpreted as replay sequenceness.

To assess statistical significance, non-parametric permutation tests were employed. Hypothesised transition matrices were shuffled 1,000 times by randomly reassigning ones and zeros, and the prediction of the shuffled matrices to the empirical transition matrix was repeated, yielding a null distribution of replay sequenceness values. Replay sequenceness exceeding the 95th percentile of this null distribution was considered significant. Candidate time lags ranging from 10 ms to 600 ms in 10 ms increments were evaluated to identify the optimal replay speed, with corrections applied for multiple comparisons across time lags.

### Resting-state network modelling

Resting-state networks were characterised using a time-delay embedded hidden Markov model (TDE-HMM) approach (Gohil et al., 2024; Higgins et al., 2021; Vidaurre et al., 2018). This method models brain activity as a sequence of latent states, each associated with distinct spectral and spatial properties.

At each time point 𝑡, the observed data 𝑌*_t_* are assumed to be generated by a latent state 𝜃*_t_*, with the probability 𝑝(𝑌*_t_* ∣ 𝜃*_t_*). The latent states evolve according to a Markov process, with transition probabilities 𝑝(𝜃*_t_* ∣ 𝜃*_t_*_-1_), which describe state transitions. This differs from the transition matrix used for replay quantification, which models reactivation probabilities between objects. The observation model uses time-embedded data vectors 𝑣𝑒𝑐(𝑌*_t_*_-1:*t*+1_) , where 𝑙 defines the embedding window (set to 𝑙 = 7), and 𝑣𝑒𝑐 denotes vectorisation. Dimensionality reduction via principal component analysis (PCA) is applied for computational efficiency, retaining the top 80 components. Each state’s observation model is characterised by a multivariate normal distribution with state-specific mean *𝑚_k_* and covariance 𝐶*_k_* , capturing both power spectra and coherence patterns, with equation (3).

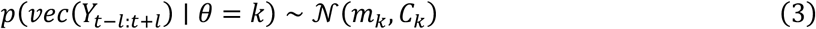

Bayesian inference with stochastic gradient variational Bayes was employed to estimate model parameters, optimising both the observation model and transition probabilities. The model was trained five times with random initialisations to avoid local minima, selecting the solution with the lowest free energy.

The trained model provided activation probability time series for 12 RSN states, their spectral properties, and the transition probability matrix. RSN states were reordered based on minimising distances between states with high transition probabilities. The most probable sequence of RSN states (Viterbi path) was computed, assuming mutual exclusivity of states.

Temporal properties such as fractional occupancy, mean interval time, mean lifetime, switching rate, and Fano factor were calculated for each RSN state. Spectral analyses involved computing mean power and coherence across frequencies, with spatial distributions visualised for each state (**Supplementary Figure 4A**).

### Replay-aligned RSN-state activation

To investigate whether replay events coincided with RSN-state activation, replay onset times were identified, and RSN-state activation around these events was analysed.

Replay onsets were defined as times when the strong reactivation of one object was immediately followed by the strong reactivation of the subsequent object in the sequence. Among all candidate time lags, the one yielding the highest replay sequenceness was selected as the optimal time lag. Using the identified time lag, we constructed the reactivation matrix 𝑋 and its time-shifted version 𝑋(𝛥𝑡). The original reactivation matrix 𝑋 was then multiplied by the hypothesised transition matrix *𝑇_r_*, representing the expected unscrambled sequences. Next, the 𝑋 × 𝑇_!_ was element-wise multiplied with the time-shifted matrix 𝑋(𝛥𝑡), yielding a matrix with each column corresponding to an object and each row representing a time point in the resting-state data. To capture the strength of replay over time, we summed the values across columns, producing a long vector 𝑅, with each element indicating the replay strength at a specific moment. We then applied a threshold to 𝑅, selecting only values above the 99th percentile to define significant replay moments. To further ensure temporal independence between replay events, we imposed a minimum gap between consecutive replay onsets, ensuring that each onset was preceded by at least 100 ms without replay activity. Specifically,

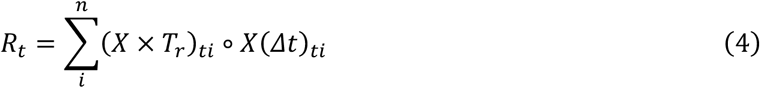

Using the identified replay onsets and the activation probability time series for each RSN state, replay-aligned RSN-state activation was analysed at the participant level. For each participant, the activation probability time series of each RSN state was baseline-corrected by subtracting the average activation probability across all time points within the session. This ensured that positive values reflected higher-than-average activation probabilities, while negative values indicated lower-than-average activation probabilities. The corrected time series was segmented into epochs centred around replay events, with a window spanning from 0.5 s before to 0.5 s after each event (0.2 s before to 0.3 s after for visualisations in **Figure 4**). The RSN-state activation probabilities were averaged across these epochs for each participant. The mean of these averaged values across participants was computed to quantify the expected increase or decrease in RSN-state activation probability in response to replay events. To assess the statistical significance of replay-aligned RSN-state activation, a permutation-based approach was employed. Specifically, the sign of the activation probabilities was randomly flipped across trials 5,000 times to generate a null distribution. The observed mean activation probability was then compared against this null distribution to determine whether it exceeded the chance level. Statistical significance was set at *p* < 0.001.

### Grid-like code

We applied representation similarity analysis (RSA) to examine grid-like neural representations (Bellmund et al., 2016; Kriegeskorte, 2008; the method used in this study is detailed in Qu et al. (2025). First, we built a model representational dissimilarity matrix (RDM) based on the angular differences between inferred directions, assuming that grid-like patterns would show greater similarity at specific angles (0–15° and 45– 60°) due to the periodic nature of grid codes. Pairs of directions within these angles were considered similar (assigned a value of zero), while other pairs were marked as dissimilar (assigned a value of one). Next, we ran a whole-brain searchlight analysis. For each voxel, we estimated activity patterns by modelling separate regressors for each direction during the GLM analysis. We then calculated a neural RDM within a searchlight sphere around each voxel by measuring correlation distances between these patterns. Finally, we compared the model RDM to the neural RDM using Pearson’s correlation, transformed these correlation values to *z*-scores, and mapped them back to the central voxel of each sphere for further analysis.

### Theta phase locking

To investigate theta phase locking of replay events and its relationship with age and DMN properties, we extracted theta phases at replay onset from the source-space MEG data for each ROI. Using non-negative matrix factorisation (NNMF), we separated the frequency range of 1–45 Hz into four distinct bands, as illustrated in **Supplementary Figure 4D**. The theta band was defined as 2–6 Hz. The data utilised for constructing the HHM were filtered to the theta band and subsequently Hilbert transformed to obtain a time series of phases.

Source-space MEG data can exhibit sign ambiguity, resulting in uncertainty in phase estimation (Vidaurre et al., 2016). To mitigate this issue, we applied a sign-flipping procedure based on the premise that high gamma power is greater at the theta trough than at the theta peak (Canolty et al., 2006). The high gamma band was defined as 80–120 Hz, corresponding to observed ripple band power increases (**Figure 2H**) and consistent with previous studies (Heusser et al., 2016). Following phase adjustment, we analysed the theta phases at replay onset.

A potential confound is that the global theta signal, representing activity across sensor space, could artificially inflate decoding accuracy at the theta peak and trough (Vidaurre et al., 2021). This might falsely suggest that replay events cluster around these phases even without true alignment. To address this, we conducted a control analysis by shuffling the decoder weights 1,000 times and using the shuffled decoders to decode the original resting-state MEG data. The resulting reactivation time series from the shuffled decoders were then used to identify replay onsets. This approach effectively disrupted the relationship between replay events and the true data while preserving the statistical structure of original signal. Replay onsets identified from the shuffled data did not show significant phase alignment with age, confirming that the observed replay onset phase alignment is not driven by global theta signal effects.

### Event-related phase-amplitude coupling

Phase-amplitude coupling (PAC) was assessed to explore interactions between theta phase and gamma amplitude during replay events. PAC was computed using the Gaussian-Copula Mutual Information (GCMI) method (Combrisson et al., 2020; Ince et al., 2017), a non-parametric approach estimating mutual information between phase and amplitude components while accounting for their marginal distributions.

High-frequency gamma amplitudes (80–120 Hz) were extracted using bandpass filtering and the Hilbert transform. Theta phases (2–6 Hz) were obtained using overlapping 1 Hz steps to minimise frequency-specific artefacts. PAC values were calculated for each trial and participant.

Statistical significance was determined using an amplitude-shuffling procedure, generating a null distribution of PAC values by randomly reordering amplitude time series 200 times per trial. Observed PAC values were converted to *z*-scores, and group-level *t*-tests assessed significance across participants. Regions showing significant PAC (*p* < 0.05) were identified.

### Connectome-based predictive modelling

Connectome-based predictive modelling (Scheinost et al., 2019; Shen et al., 2017) was employed to assess whether resting-state functional connectivity within the DMN could predict task-induced 2D replay sequenceness.

Using preprocessed fMRI data, BOLD time series from 400 brain regions (including 91 regions within the DMN) were extracted based on the Schaefer atlas (Schaefer et al., 2018). Pearson correlations between time series of each pair of regions were calculated and Fisher *z*-transformed to create individual rsFC matrices.

Leave-one-out cross-validation (LOOCV) was used to train and validate the predictive model. In each iteration, data from one participant were held out as the test set, and the remaining data formed the training set. Feature selection within the training set identified rsFC edges significantly correlated with replay sequenceness (*p* < 0.001). The sum of strengths of positively and negatively correlated edges served as predictors in a linear regression model.

Predicted replay sequenceness values were generated for each left-out participant. Model performance was evaluated using Spearman’s correlation between predicted and actual replay sequenceness. Permutation tests with 10,000 iterations assessed the statistical significance by comparing model performance to a null distribution generated by shuffling replay sequenceness values. Statistical significance was established if the *p*-value from the permutation tests was less than 0.05.

### Mediation analysis

To investigate whether DMN rsFC mediated the relationships between age, task-induced replay sequenceness, and inference efficiency, mediation analyses were conducted following standard procedures (Preacher & Hayes, 2004). The analysis aimed to determine whether the effect of the independent variable (e.g., age) on the dependent variable (e.g., replay sequenceness or inference efficiency) could be explained, in whole or in part, by the mediator variable (e.g., DMN rsFC).

Bootstrapping methods with 5,000 resamples were employed to generate 95% bias-corrected confidence intervals for the indirect effects, allowing for robust estimation of mediation effects without reliance on normality assumptions. All variables were mean-centred prior to analysis to facilitate interpretation, and standardised coefficients were reported to enable comparison across variables. An indirect effect was considered significant if its confidence interval did not include zero, indicating a reliable mediation effect. Statistical significance for individual paths in the mediation model was determined at the *p* < 0.05 level.

### Statistical analysis

Statistical analyses were conducted using Python, employing the scipy.stats and statsmodels libraries for hypothesis testing and statistical modelling. Throughout the study, two-tailed *p*-values were used. Corrections for multiple comparisons were applied where appropriate to control the family-wise error rate, particularly in analyses involving multiple time points or ROIs.

Correlation analyses were performed using Pearson’s correlation coefficient (*r*) for normally distributed data and Spearman’s rank correlation coefficient (*ρ*) for non-parametric data or when assessing monotonic relationships. These analyses examined relationships between variables such as age, decoding accuracy, replay sequenceness, and DMN rsFC.

For replay detection, non-parametric permutation tests were employed to assess the statistical significance of replay sequenceness. Hypothesised transition matrices were shuffled 1,000 times to generate null distributions, and corrections for multiple comparisons were applied across the range of time lags assessed (10 ms to 600 ms in 10 ms increments). Replay sequenceness exceeding the 95th percentile of the maximum values from the null distributions was considered statistically significant.

To evaluate replay-aligned RSN-state activation and phase-amplitude coupling, permutation-based approaches were used. For RSN-state activation, the sign of activation probabilities was randomly flipped across trials 5,000 times to create a null distribution, and statistical significance was set at *p* < 0.001. For PAC analysis, an amplitude-shuffling procedure with 200 repetitions per trial generated null distributions, and observed PAC values were converted to *z*-scores for statistical testing. Within-subject and group-level statistics were calculated, with significance determined at *p* < 0.05.

Circular statistics were employed for analyses involving angular data, such as theta phase locking. The Rayleigh test was used as a one-sample test for non-uniformity of circular data, assessing whether replay events occurred preferentially at specific phases. The Watson-Williams test served as a circular analogue of the two-sample *t*-test, comparing mean phases between age groups. Circular-linear correlation coefficients were calculated to assess relationships between angular variables and linear variables like age.

In the connectome-based predictive modelling, model performance was evaluated using Spearman’s correlation between predicted and actual values of task-induced replay sequenceness. Permutation tests with 10,000 iterations assessed the robustness of the predictive models by comparing the observed performance to that expected by chance. Statistical significance was established if the *p*-value from the permutation tests was less than 0.05.

For mediation analyses, bootstrapping methods with 5,000 resamples were used to estimate indirect effects and generate 95% bias-corrected confidence intervals. An indirect effect was considered significant if the confidence interval did not include zero. Variables were mean-centred, and standardised coefficients were reported to facilitate comparison across variables.

To compare the strength of two dependent correlation coefficients (i.e., correlations sharing one variable), Steiger’s *Z*-test was used. This test assessed whether the difference between two correlations was statistically significant. For example, between age and replay–DMN alignment versus age and replay–VIS alignment. All comparisons were two-tailed, and statistical significance was set at *p* < 0.05.

## Acknowledgment

Conceptualisation, Y.L., Y.Q., L.P., and T.B.; Investigation, L.P., Y.Q., J.O., S.W., Y.L., C.G., M.W, and T.B.; Writing — Original Draft, L.P., Y.L.; Writing — Review & Editing, Y.L., L.P., Y.Q., T.B. This study is supported by the National Science and Technology Innovation 2030 Major Program (2022ZD0205500), the National Natural Science Foundation of China (32271093), the Beijing Natural Science Foundation (Z230010, L222033), and the Fundamental Research Funds for the Central Universities.

## Data availability

MEG, fMRI and behavioural data that support the conclusions in this study will be available on http://zenodo.org/uploads/14177579 upon publication.

## Code availability

The analysis code will be publicly available on https://gitlab.com/liu_lab/replay_development upon publication.

## Supplementary Information

**Supplementary Figure 1.**
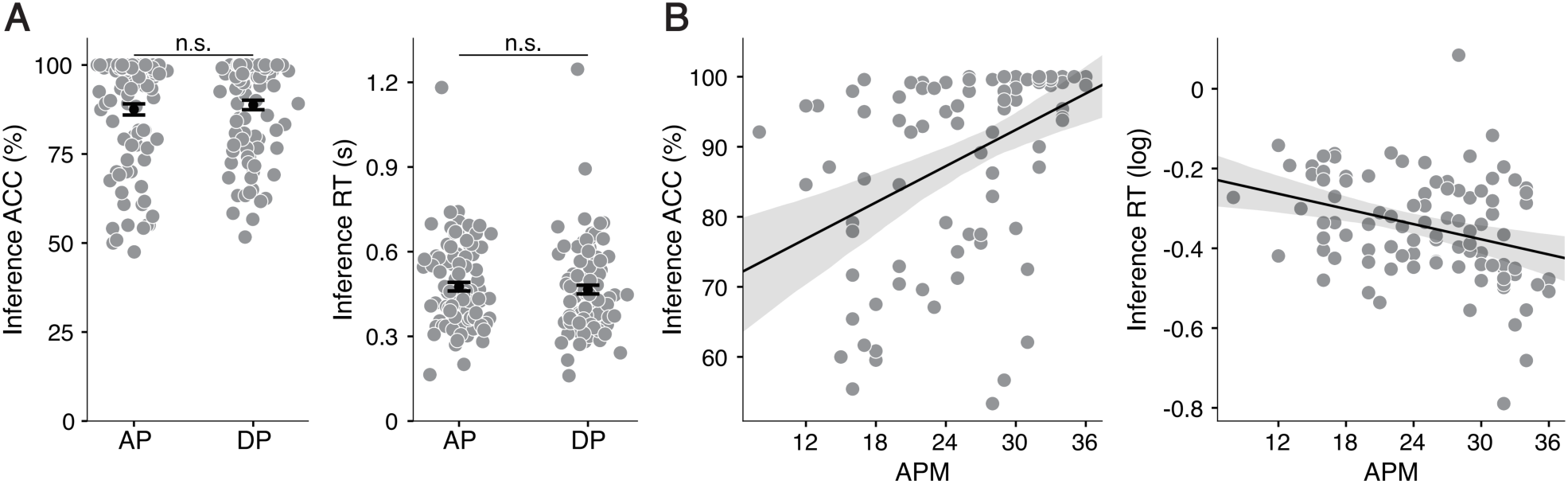
Task performances in map-based inference and general reasoning ability. **(A)** Accuracy (left) and reaction time (right) in the map-based inference task showed no difference between the attack (AP) and defence (DP) dimensions. Error bars represent the group SEM. ‘ns’ for non-significant. **(B)** Relationship between reasoning ability and map-based inference task performance: Accuracy positively correlated with the Advanced Progressive Matrices (APM) score (left), while reaction time negatively correlated with the APM score (right). The solid line represents the best-fitting line across participants, and the shaded area indicates the 95% confidence interval. Each dot represents an individual participant.

**Supplementary Figure 2.**
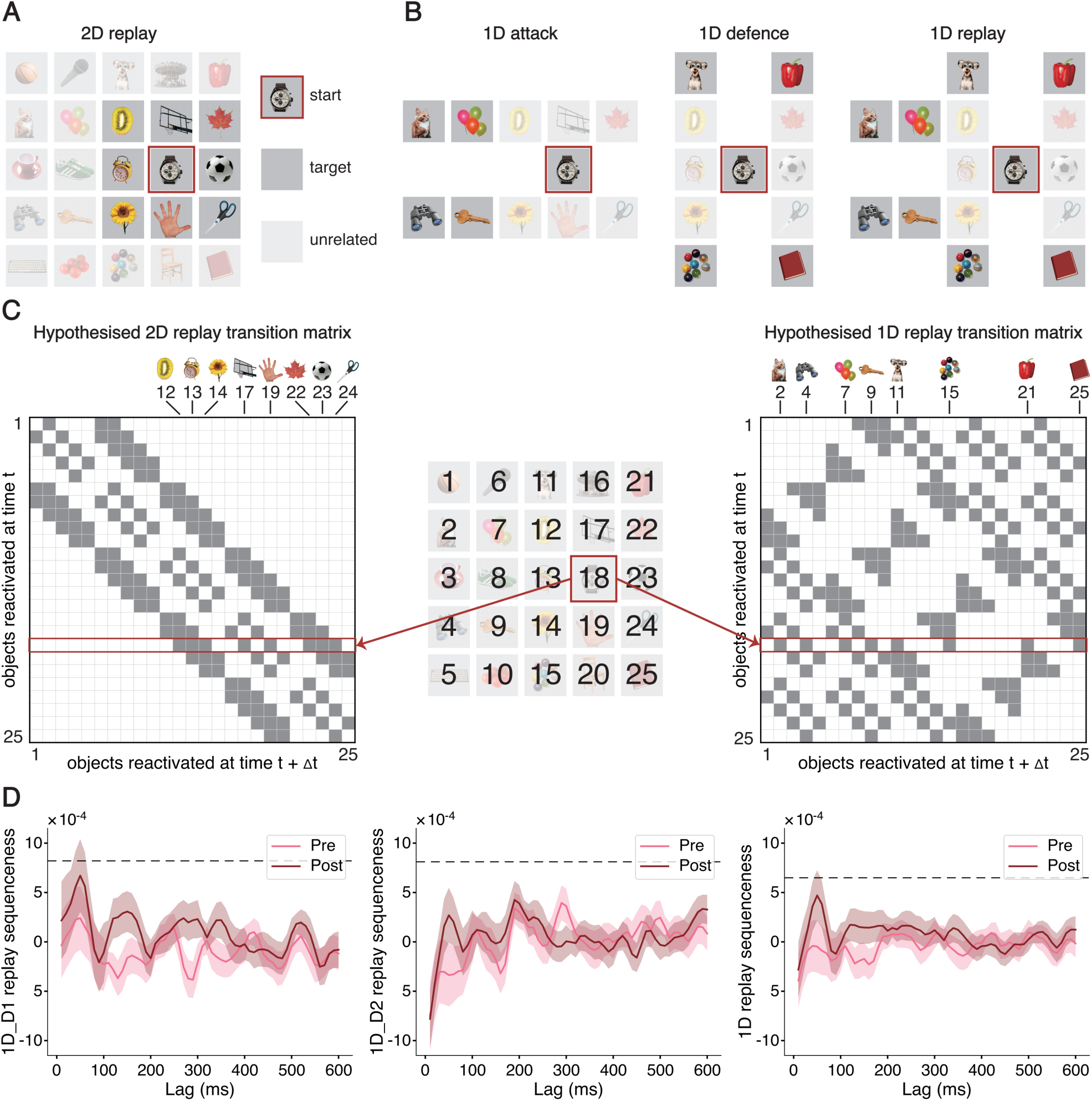
No 1D replay detected in post-task rest. **(A)** An illustration of 2D replay. 2D replay starts from the object in the red frame and targets the blacked objects. In 2D replay, these targets are adjacent to the starting object in the 2D space formed by the AP and DP dimensions. **(B)** Illustrations of three types of 1D replay: 1D replay of the attack power dimension (left), defence power dimension (middle), and the combination of both (right). In 1D replay, target objects are one level away from the starting object in either the AP or DP dimension, excluding stimuli that are neighbours in the 2D map, to distinguish from 2D replay. **(C)** Hypothesised transition matrices. Hypothesised 2D (left) and 1D (right) replay transition matrices were derived by unfolding the 5 × 5 map (middle). Gray cells indicate transferable transitions; white cells indicate non-transferable transitions. The highlighted rows illustrate transitions of the starting object “watch”. **(D)** Unlike 2D replay, replay sequenceness for all types of 1D replay fell below the permutation threshold (dashed black line). No replay of 1D was observed. The shaded area represents the group SEM.

**Supplementary Figure 3.**
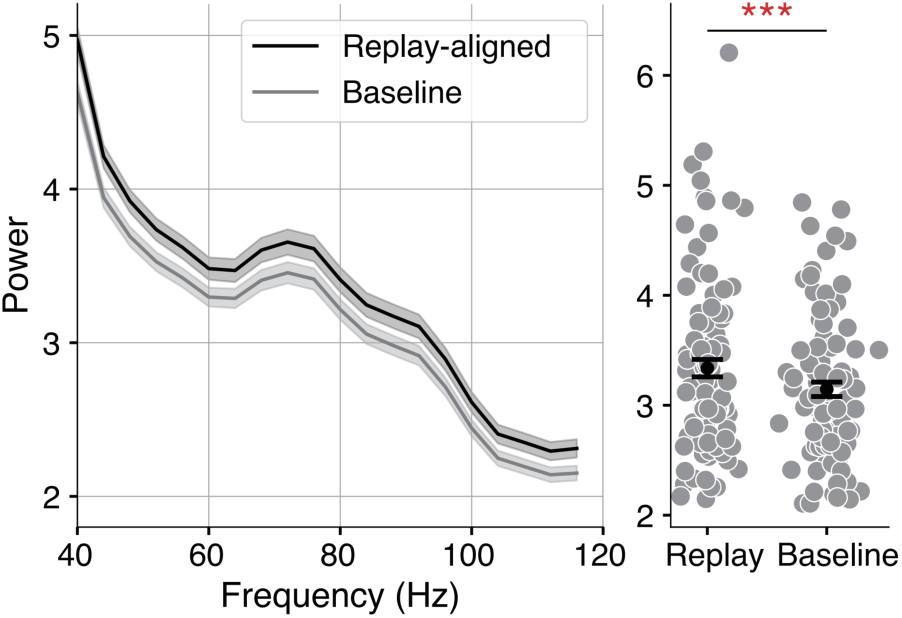
Increased high-frequency band power around replay events. High-frequency power increase associated with replay events. Left: The onset of replay events is associated with an increase in high-frequency power relative to the global average (mean ± SEM over participants). Right: Replay-aligned power was computed by taking 100 ms windows of data around identified replay onsets. The averaged replay-aligned power in the 80–120 Hz band is significantly higher than the baseline. ‘***’ for *p* < 0.001.

**Supplementary Figure 4.**
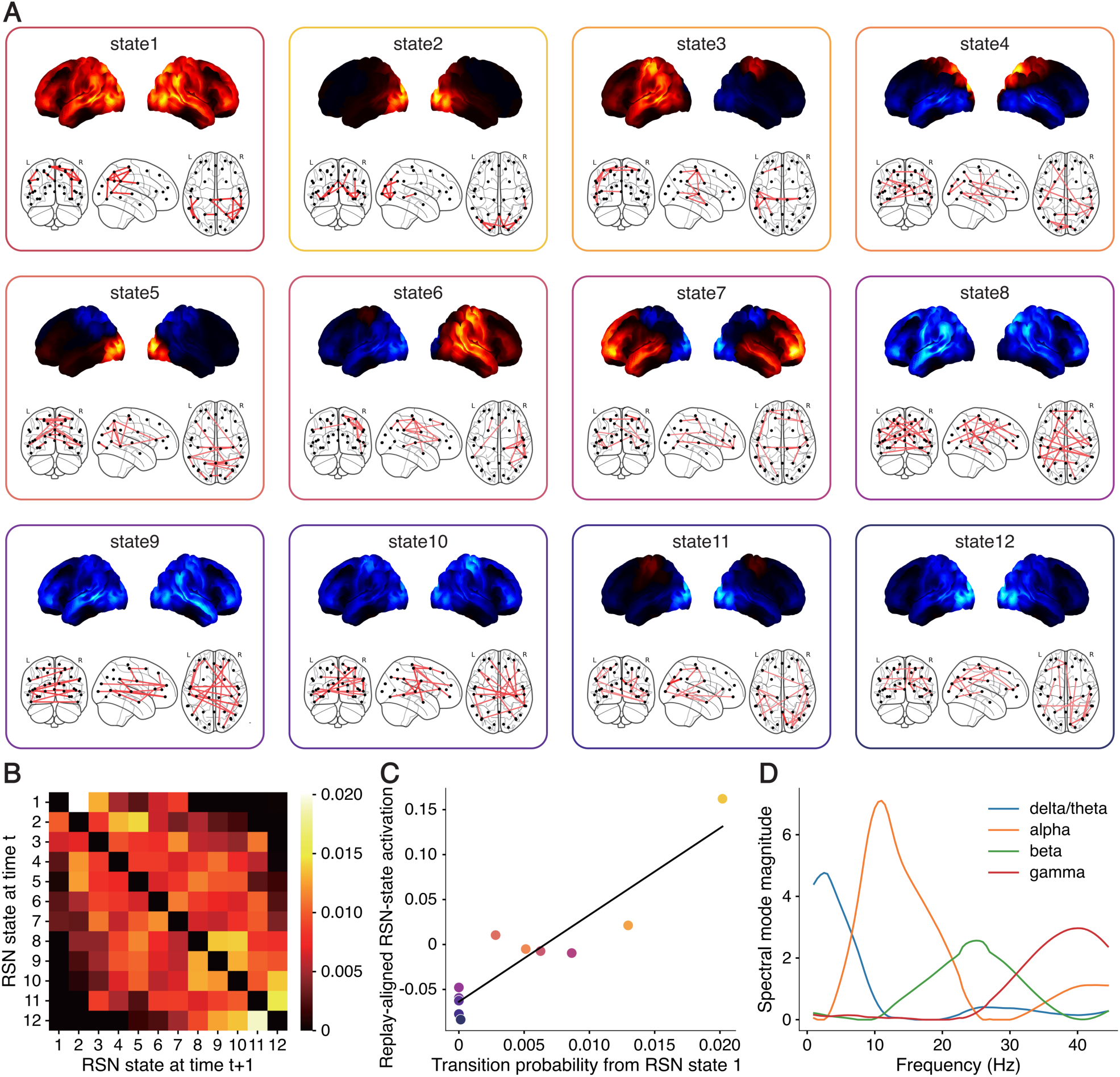
The broadband power and phase-locking network properties of all RSN states. **(A)** Visualisation of RSN states. Twelve RSN states were identified from the post-task resting MEG time series, using the HMM with time-delayed embedding (Vidaurre et al., 2018). Each RSN state corresponds to a distinct power distribution and phase-locking pattern between ROIs. The power maps (sagittal view) and the phase-locking maps of each RSN state are plotted in each cell. The power maps were normalised across all states by subtracting the mean, and the phase-locking maps were thresholded using a Gaussian mixture model. **(B)** RSN-state transition probability matrix. This matrix illustrates the likelihood of transitioning from one state at time 𝑡 (horizontal axis) to another state at time 𝑡 + 1 (vertical axis). Each entry represents a specific transition probability. **(C)** Replay-aligned RSN-state activation positively correlated with its transition probability from the DMN for all other RSN states. **(D)** Spectral factorisation. Using non-negative matrix factorisation, the 1–45 Hz frequency range was decomposed into four bands: theta (2–6 Hz), alpha (6–12 Hz), beta (12–32 Hz), and gamma (32–45 Hz).

**Supplementary Figure 5.**
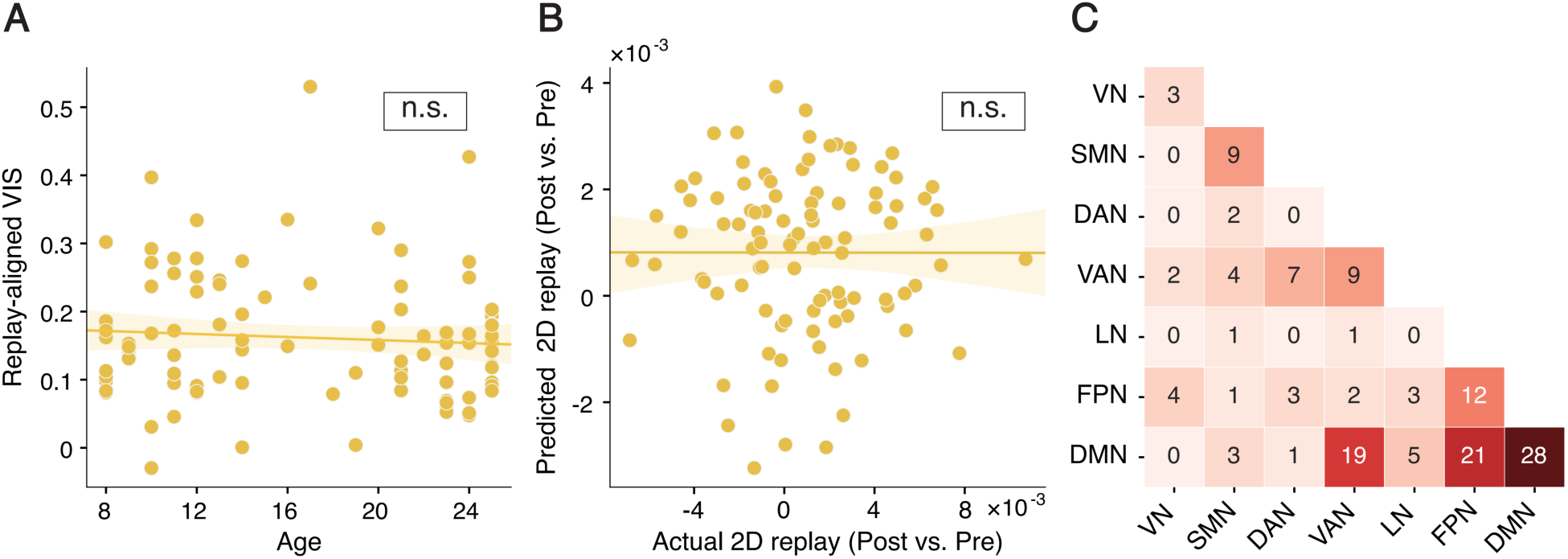
Control analysis on the relationships between the VIS and replay. **(A)** In contrast to the DMN, replay-aligned VIS activation was not correlated with age. ‘ns’ for non-significant. **(B)** Unlike the DMN, the VIS rsFC failed to predict replay, showing no significant correlation between predicted and actual replay sequenceness. The solid line represents the best-fitting line across participants, and the shaded area indicates the 95% confidence interval. Each dot represents an individual participant. ‘ns’ for non-significant. **(C)** Distribution of robust features in the valid predictive model using whole-brain rsFC, with features within and across the DMN contributing more to the prediction.

**Supplementary Figure 6.**
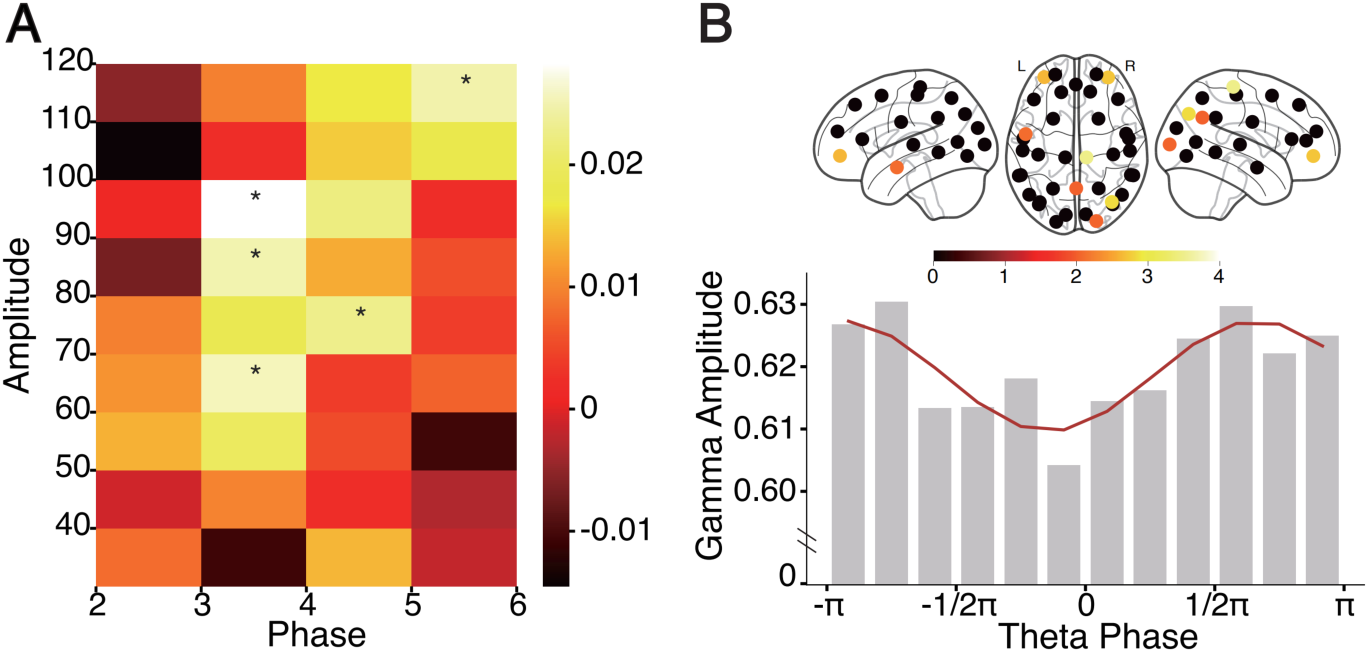
Event-related phase-amplitude coupling of replay. **(A)** ERPAC analysis in the PCC. Event-related phase-amplitude coupling (ERPAC) of replay events was analysed in the PCC. The heat map shows Gaussian-Copula PAC values (Combrisson et al., 2020; Ince et al., 2017), across theta phases and high-frequency amplitudes (see Methods for details). A significant coupling between theta phase and gamma amplitude is observed. ‘*’ for *p* < 0.05. **(B)** Spatial distribution and phase dependency. Top: Spatial distribution of ROIs, with regions showing significant ERPAC of theta phase and gamma amplitude coloured according to their t-test values at the group level. Bottom: Gamma power changes with theta phase in the PCC around replay events, showing higher amplitude at the theta trough compared to the theta peak.

**Supplementary Figure 7.**
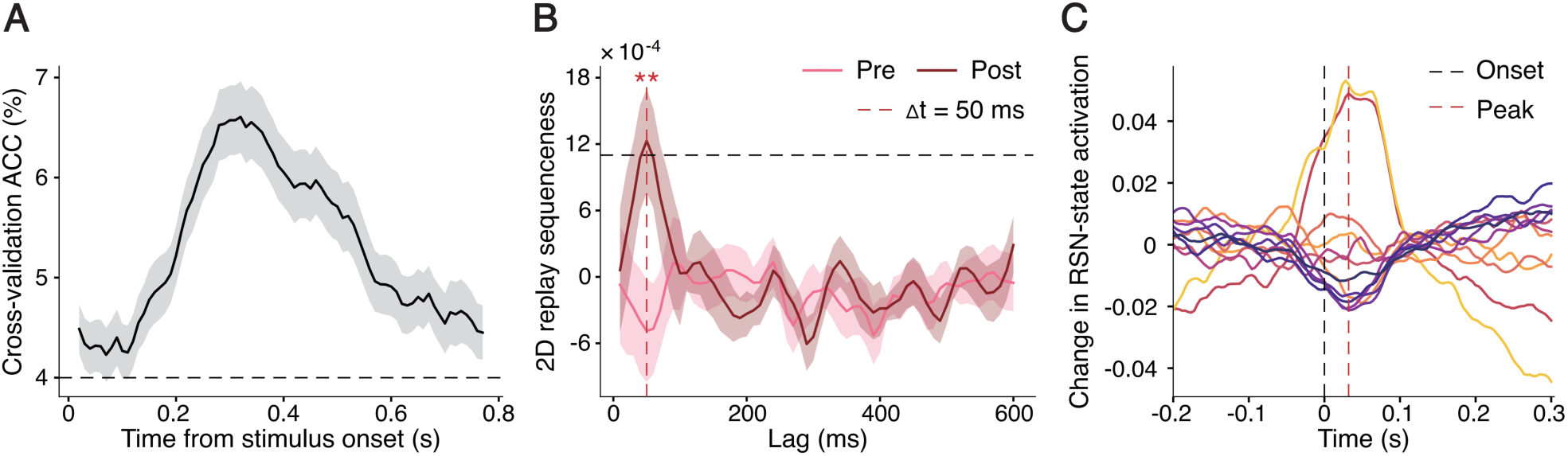
Replication of main effects in younger participants. **(A)** Decoding accuracy over time. Cross-validation results revealed that peak decoding accuracy was achieved at 300 ms post-stimulus onset, consistent with findings in the full sample. Error bars represent the group SEM. **(B)** 2D replay during rest. 2D replay during post-task rest was significantly higher than expected by chance (black dashed line indicates the permutation threshold) at time lags of 50–70 ms. Additionally, post-task replay was significantly higher than pre-task replay at the 50 ms lag, replicating the replay effects observed in the full sample. Error bars represent the group SEM. ‘**’ for *p* < 0.01. **(C)** Changes in RSN-state activation around replay events. Replay events accompanied by a transient increase in RSN-state activation, particularly in the DMN and VIS, replicating the temporal profile observed in the full sample. This increase was observed from −0.2 s to 0.3 s relative to replay onset (dashed black line), peaking around 30 ms after replay onset (dashed red line).

**Supplementary Figure 8.**
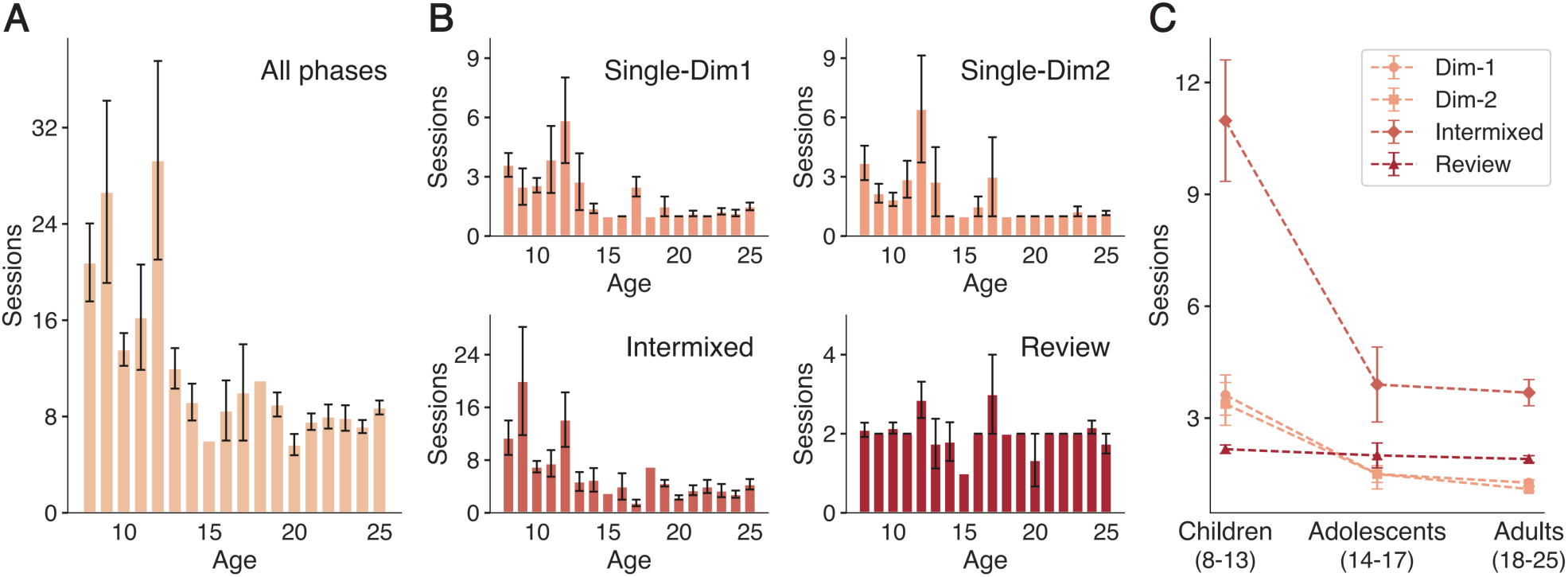
Distribution of training sessions across age groups. **(A)** Total number of training sessions required to reach memory criterion (accuracy ≥ 80%) across all phases, plotted by age (8–25 years). Error bars represent the group SEM. **(B)** Number of sessions completed in each training phase: single-dimension training for dimension 1 and dimension 2, intermixed training, and final review. Error bars represent the group SEM. **(C)** Interaction between age groups (children: 8–13; adolescents: 14–17; adults: 18–25) and phases, indicating that children struggled most during the intermixed phase. Error bars represent the group SEM.

**Supplementary Figure 9.**
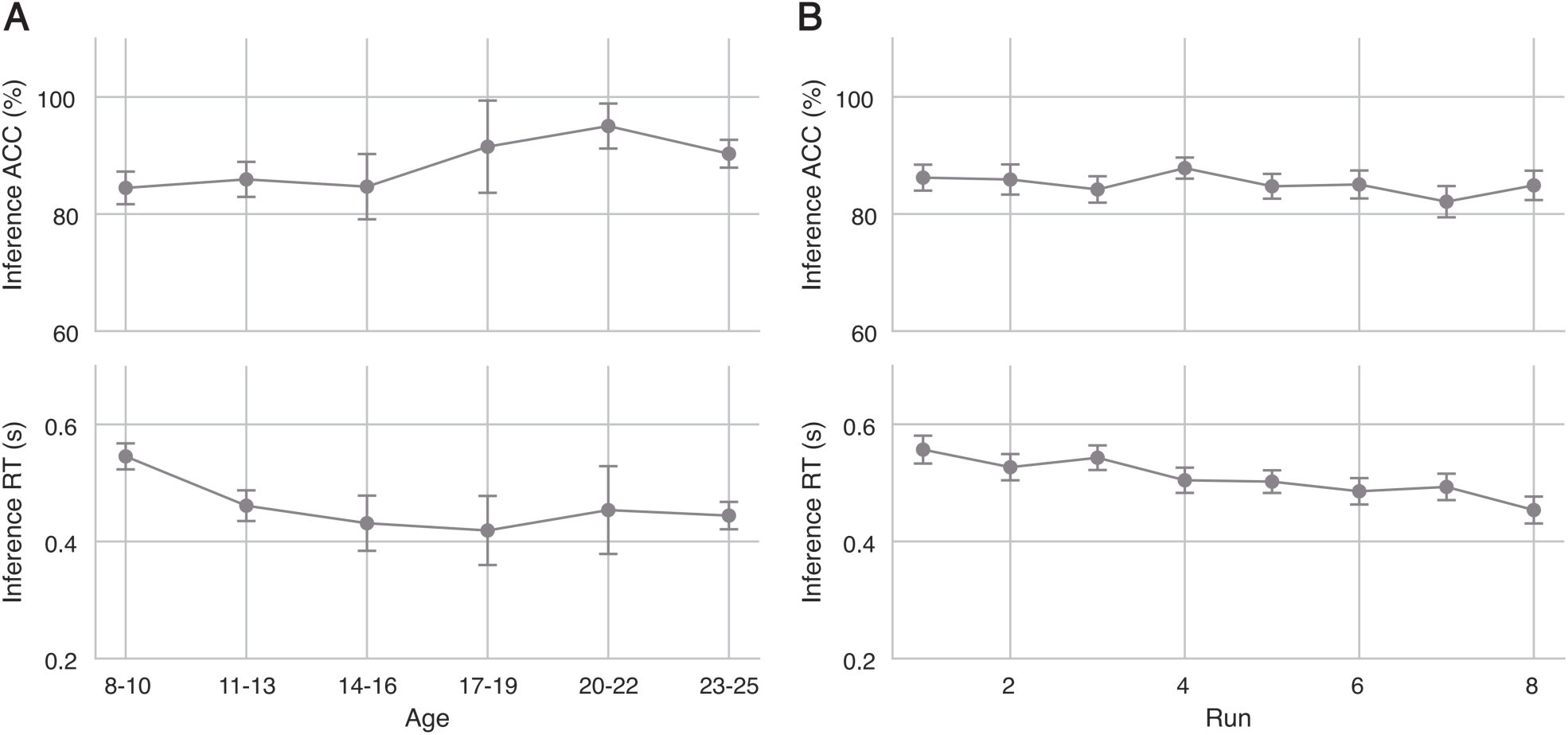
Age-related and temporal stability of behavioural performance. **(A)** Behavioural performance across age groups. Behavioural performance in the inference task was analysed across different age groups. All groups performed significantly above chance level, demonstrating active task engagement. Error bars represent the group SEM. **(B)** Performance stability over task runs. Behavioural performance of participants aged 8–13 was examined across runs of the inference task. Accuracy remained stable over time, while reaction time showed changes, suggesting possible familiarity effects rather than fatigue. Error bars represent the group SEM.

**Supplementary Figure 10.**
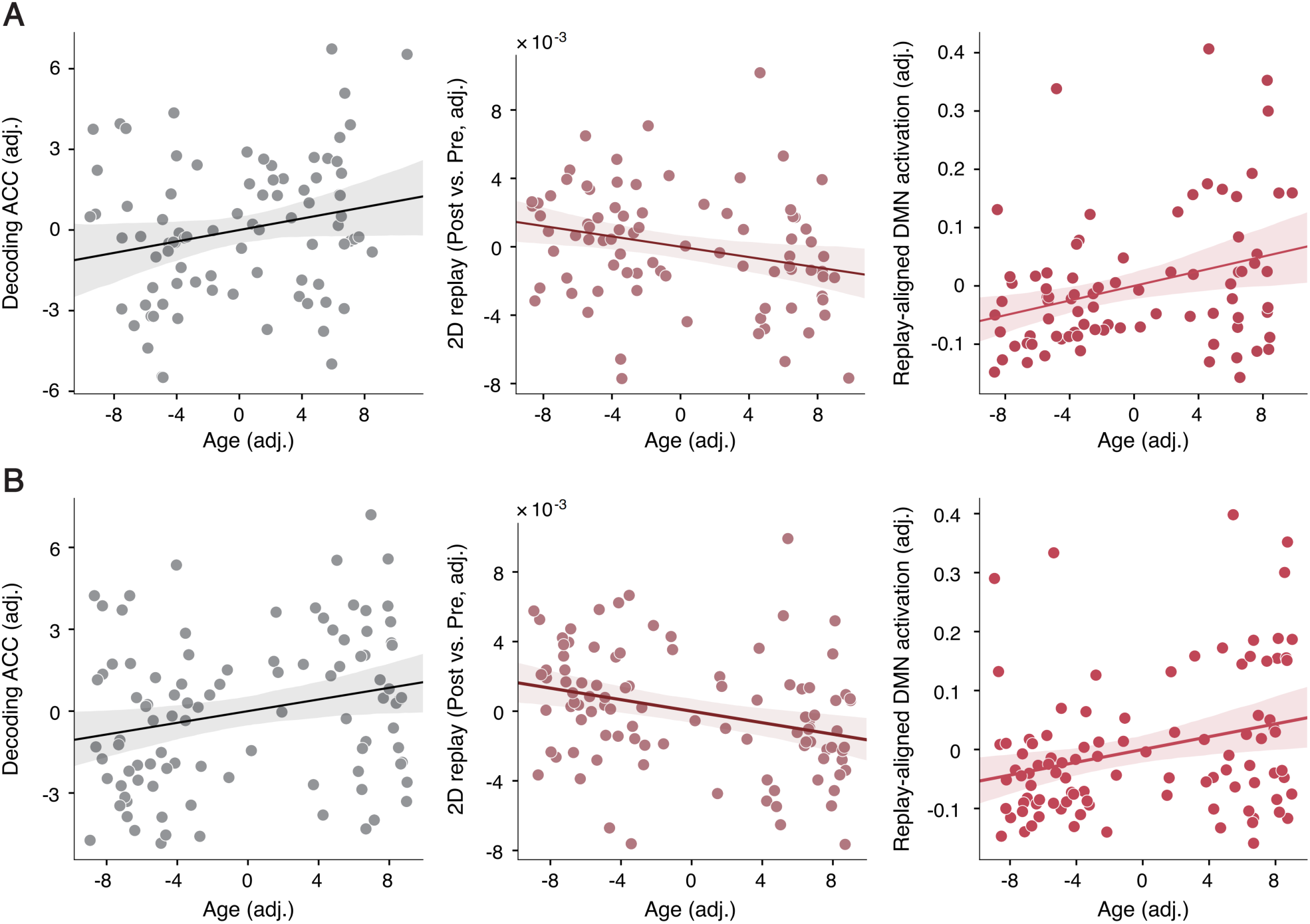
Control analyses of head motion and head size. **(A)** Controlling for head motion. The correlations between age and decoding accuracy (left), task-induced 2D replay sequenceness (middle), and replay-aligned DMN activation (right) remained significant after controlling for mean head motion during the corresponding MEG scanning sessions. **(B)** Controlling for head size effects. The correlations between age and decoding accuracy (left), task-induced 2D replay sequenceness (middle), and replay-aligned DMN activation (right) remained significant after controlling for the mean lead field norms.

## Supplementary Note 1. Control analyses of age-related confounding factors

To ensure the robustness of our findings, we conducted comprehensive control analyses to account for potential age-related confounding factors.

### Main effects across age group

We assessed whether the main effects of replay and DMN activation were influenced by participant age by dividing participants into younger (age < 15) and older (age ≥ 15) groups. For pre-task and post-task 1D replay, as well as pre-task 2D replay, no significant effects were observed in either group. Post-task 2D replay showed significant results in both groups. Notably, the increase in 2D replay from pre-task to post-task was significant also in the younger group (paired *t*-test, *p* < 0.01) but not in the older group. This supports our interpretation that younger participants required more replay to consolidate the cognitive map. The general properties of the DMN and replay-aligned DMN activation were consistently replicated across both age groups. **Supplementary Figure 7** illustrates that the observed effects are not solely driven by older participants.

### Training procedure

All participants completed a three-stage training procedure prior to the formal experiment to ensure full memorisation of 200 pairwise relationships, as described in Qu et al. (2025). The first stage was single dimension learning, participants learned one-rank relationships separately for each dimension until reaching at least 80% accuracy. The second was intermixed learning, trials from both dimensions were intermixed, and incorrectly recalled pairs were relearned until performance again exceeded 80%. In the final review stage, all learned pairs were revisited across three days to stabilise memory before the formal experiment. See **Supplementary Figure 8** and **Supplementary Table 4** for training sessions across age groups and training phases. This adaptive procedure accommodated age-related differences in learning speed while enforcing a uniform memory accuracy criterion. Although the number of training sessions required to reach criterion was significantly correlated with age, *r* = −0.483, *p* < 0.001, memory performance after training did not show a significant association with the number of training sessions, *r* = −0.087, *p* = 0.405, nor with age, *r* = 0.074, *p* = 0.471. Moreover, all main correlations reported in the present study remained significant after controlling for individual differences in memory performance, suggesting that the observed age-related changes were not driven by differences in memory accuracy.

### Task engagement

To ensure active task engagement, we set a stringent pre-scanning memory criterion, allowing participants to proceed to scanning sessions only after achieving at least 80% accuracy in pairwise comparisons during training. Inference accuracy for all participants exceeded the 50% chance level. **Supplementary Figure 9A** shows behavioural performance across age groups, confirming that all groups performed significantly above chance (one-sample *t*-tests, *p* < 0.001). Children aged 8–13 performed comparably to other age groups, indicating effective task engagement.

### Fatigue effect

We examined potential fatigue effects in youngest children aged 8–13 by analysing behavioural performance across the eight runs of the inference task (**Supplementary Figure 9B**). Repeated measures ANOVA revealed no significant differences in accuracy across runs (*F*(7, 294) = 1.796, *p* = 0.112), indicating stable performance.

Reaction time showed significant changes across runs (*F*(7, 294) = 5.868, *p* < 0.001), suggesting familiarity effects rather than fatigue.

### Head motion

Head motion is a common concern in developmental neuroimaging studies (Wehner et al., 2008). We calculated mean head motion for each participant during task and rest periods. After controlling for head motion, the correlations between age and decoding accuracy (partial *r* = 0.220, *p* = 0.043), task-induced 2D replay (partial *r* = −0.261, *p* = 0.017), and replay-aligned DMN activation (partial *r* = 0.316, *p* = 0.004) remained significant (**Supplementary Figure 10A**). This indicates that head motion did not confound our results.

### Head size effects

Head size is a known confound in neurodevelopmental MEG studies due to the fixed sensor geometry of the MEG helmet (Johnson & He, 2019), which can result in smaller brains being farther from the sensors, leading to lower signal strength and SNR in younger participants (Brookes et al., 2018). To assess the potential impact of head size on our results, we followed Brookes et al. (2018), and simulated lead fields for a single source in 38 cortical regions for each participant and examined the correlation between lead field norms (as a proxy for signal strength) and age. No significant linear or non-linear relationship was found between the averaged lead field norms and age (all *p* > 0.1), and the inclusion of lead field norms as a regressor of no interest did not alter the primary results (**Supplementary Figure 10B**). This indicates that head size had no measurable confounding effect on our analyses.

### Processing speed

To ensure that younger participants were not disadvantaged by processing speed, we set a decision time limit of 2 seconds, based on pilot testing with children and adults. Behavioural data confirmed that all participants responded well within this limit (one-sample *t*-test against 2 seconds: *t*(97) = −102.630, *p* < 0.001), including children aged 8–13 (*t*(42) = −81.179, *p* < 0.001). Thus, differences in processing speed did not account for our findings.

### Non-linear age-related development hypothesis

As suggested in Brookes et al. (2018), we compared three candidate models to test whether nonlinear models provided a better fit for variables significantly correlated with age: a linear model 𝑦 = 𝑎_1_𝑥 + 𝑎_2_; a non-linear monotonic model 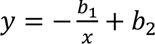; a quadratic model 𝑦 = 𝑐_1_𝑥^2^ + 𝑐_2_𝑥 + 𝑐_3_. The linear models of inference accuracy, decoding accuracy, task-induced 2D replay, replay-aligned DMN activation and grid-like code consistently outperformed the non-linear and quadratic alternatives, as indicated by higher adjusted R-squared values and lower Bayesian Information Criterion (BIC) scores. For temporal and spectral characteristics of the DMN that did not show a significant linear association with age, neither the non-linear monotonic model nor the quadratic model provided a significant improvement in fit. These results suggest that linear relationships adequately describe our data, with no substantial evidence supporting nonlinear age effects.

### Socioeconomic status (SES)

Due to privacy concerns, valid SES data were available for only 51 participants. Analysis revealed a significant negative correlation between age and SES (Spearman’s *ρ* = −0.326, *p* = 0.019), indicating higher SES in younger participants. However, none of the key variables in our study showed significant correlations with SES (all *p* > 0.1), suggesting that SES did not influence our results.

### Time of day

All MEG scanning sessions were conducted in the morning, between 10:00 am and 12:00 pm, to control for potential diurnal variations in brain activity and cognitive performance.

### Sleep quality

Participants were instructed to have adequate rest the night before scanning. Sleep quality was assessed using the Pittsburgh Sleep Quality Index (PSQI, Buysse et al., 1989). None of the participants scored above the cutoff of 10.

**Table S1.** Descriptive statistics of demographic variables (N = 106).

|  | Total | Group |  | t-test |
| --- | --- | --- | --- | --- |
|  |  | Age < 15<br>(N = 53) | Age ≥ 15<br>(N = 53) |  |
| **Female** | 51 | 22 | 29 |  |
| **Male** | 55 | 31 | 24 |  |
| **Age** | 16.45 ± 0.61 | 10.75 ± 0.20 | 22.15 ± 0.27 | 23.79*** |
| **APM** | 25.21 ± 0.64 | 22.10 ± 0.60 | 28.25 ± 0.54 | 5.25*** |

2 '\*\*\*' for independent-sample *t*-test $p < 0.001$ .

**Table S2.** MNI coordinates for the 38 ROIs in the source space.

| Region of interest | Hemisphere | MNI<br>coordinates |  |  | Hemisphere | MNI<br>coordinates |  |  |
| --- | --- | --- | --- | --- | --- | --- | --- | --- |
|  |  | *x* | *y* | *z* |  | *x* | *y* | *z* |
| Cuneal Cortex | L | −10 | −87 | 23 | R | 10 | −87 | 23 |
| Lateral Occipital Cortex<br>inferior division | L | −37 | −77 | −5 | R | 37 | −77 | −5 |
| Subcallosal Cortex | L | −53 | −10 | 29 | R | 54 | −10 | 29 |
| Planum Temporale | L | −54 | −22 | 6 | R | 55 | −22 | 7 |
| Postcentral Gyrus | L | −38 | −26 | 57 | R | 38 | −26 | 57 |
| Superior Parietal Lobule | L | −23 | −61 | 54 | R | 23 | −61 | 54 |
| Middle Temporal Gyrus<br>temporooccipital part | L | −47 | −67 | 9 | R | 47 | −67 | 9 |
| Lateral Occipital Cortex<br>superior division | L | −36 | −74 | 37 | R | 36 | −74 | 38 |
| Superior Temporal Gyrus<br>anterior division | L | −51 | −6 | −17 | R | 51 | −5 | −17 |
| Sensorimotor Cortex | L | −11 | −29 | 65 | R | 11 | −29 | 65 |
| Supramarginal Gyrus<br>posterior division | L | −56 | −47 | 34 | R | 56 | −47 | 34 |
| Inferior Frontal Gyrus<br>pars triangularis | L | −42 | 30 | 12 | R | 43 | 30 | 12 |
| Occipital Pole | L | −20 | −94 | 7 | R | 21 | −94 | 7 |
| Middle Frontal Gyrus | L | −23 | 10 | 56 | R | 23 | 10 | 56 |
| Superior Frontal Gyrus | L | −15 | 37 | 47 | R | 15 | 37 | 47 |
| Frontal Lobe | L | −32 | 52 | −5 | R | 33 | 52 | −5 |
| Middle Temporal Gyrus<br>temporooccipital part | L | −58 | −48 | −1 | R | 58 | −47 | −1 |
| Frontal Lobe | L | −21 | 55 | 20 | R | 21 | 55 | 20 |
| Posterior Cingulate Cortex | M | 0 | −60 | 34 |  |  |  |  |
| Medial Prefrontal Cortex | M | 0 | 45 | 10 |  |  |  |  |

**Table S3.** Pair-wise correlations across all key variables (N = 98).

|  | 1 | 2 | 3 | 4 | 5 | 6 | 7 | 8 | 9 | 10 | 11 | 12 | 13 | 14 | 15 |
| --- | --- | --- | --- | --- | --- | --- | --- | --- | --- | --- | --- | --- | --- | --- | --- |
| **1. Age** |  |  |  |  |  |  |  |  |  |  |  |  |  |  |  |
| **2. Inference ACC** | 0.219<br>* |  |  |  |  |  |  |  |  |  |  |  |  |  |  |
| **3. Inference RT (log)** | -0.259<br>** | -0.585<br>*** |  |  |  |  |  |  |  |  |  |  |  |  |  |
| **4. APM** | 0.505<br>*** | 0.416<br>*** | -0.321<br>** |  |  |  |  |  |  |  |  |  |  |  |  |
| **5. Decoding ACC** | 0.255<br>* | 0.305<br>** | -0.212<br>* | 0.263<br>* |  |  |  |  |  |  |  |  |  |  |  |
| **6. 2D replay (Pre)** | 0.167 | -0.107 | 0.017 | 0.14 | 0.025 |  |  |  |  |  |  |  |  |  |  |
| **7. 2D replay (Post)** | -0.172 | -0.11 | 0.148 | -0.075 | -0.054 | 0.370<br>*** |  |  |  |  |  |  |  |  |  |
| **8. 2D replay (Post vs. Pre)** | -0.300<br>** | -0.017 | 0.127 | -0.187 | -0.071 | -0.470<br>*** | 0.646<br>*** |  |  |  |  |  |  |  |  |
| **9. DMN occupancy** | 0.187 | 0.102 | -0.121 | 0.022 | -0.041 | -0.016 | 0.072 | 0.081 |  |  |  |  |  |  |  |
| **10. DMN interval** | -0.13 | 0.025 | 0.059 | -0.042 | 0.088 | 0.023 | 0 | -0.018 | -0.819<br>*** |  |  |  |  |  |  |
| **11. DMN power** | 0.088 | 0.127 | -0.095 | -0.005 | -0.013 | -0.009 | 0.12 | 0.122 | 0.794<br>*** | -0.607<br>*** |  |  |  |  |  |
| **12. DMN coherence** | -0.061 | 0.063 | 0.011 | -0.047 | 0.071 | -0.015 | 0.049 | 0.059 | -0.753<br>*** | 0.921<br>*** | -0.490<br>*** |  |  |  |  |
| **13. Replay-aligned DMN activation** | 0.281<br>** | 0.171 | -0.187 | 0.097 | 0.046 | -0.007 | -0.048 | -0.039 | 0.875<br>*** | -0.671<br>*** | 0.749<br>*** | -0.679<br>*** |  |  |  |
| **14. Replay-aligned VIS activation** | -0.075 | 0.155 | -0.141 | 0.077 | 0.121 | 0.202<br>* | -0.116 | -0.276<br>** | -0.171 | 0.041 | -0.068 | -0.051 | -0.042 |  |  |
| **15. fMRI DMN rsFC** | 0.398<br>*** | 0.202<br>* | -0.314<br>** | 0.278<br>** | 0.225<br>* | 0.210<br>* | -0.403<br>*** | -0.565<br>*** | -0.066 | 0.018 | -0.115 | -0.043 | 0.082 | 0.179 |  |

2 '\*\*\*' for Pearson correlation $p < 0.001$ , '\*\*' for $p < 0.01$ , and '\*' for $p < 0.05$ .

**Table S4.** Training sessions for each age group and learning phase (N = 98).

| Age group | Single-Dim | Intermixed | Review | All phases |
| --- | --- | --- | --- | --- |
| Children (8–13) | 3.50 ± 0.53 | 10.98 ± 1.61 | 2.17 ± 0.11 | 20.14 ± 2.16 |
| Adolescents (14–17) | 1.50 ± 0.28 | 3.90 ± 0.96 | 2.00 ± 0.32 | 8.90 ± 1.02 |
| Adults (18–25) | 1.18 ± 0.06 | 3.68 ± 0.34 | 1.90 ± 0.09 | 7.95 ± 0.32 |
| All participants | 2.26 ± 0.26 | 7.00 ± 0.82 | 2.03 ± 0.07 | 13.56 ± 1.15 |

2 Values are mean ± SEM. Single-Dim values are averaged across AP and DP dimensions.

